# Continuous karyotypic and cryptic structural fluctuations drive acral melanoma evolution

**DOI:** 10.64898/2026.11.18.767656

**Authors:** Eric A. Smith, Anastasia Prokofyeva, Jian Zhao, Min Hu, Rebecca G. Zitnay, Carly A. Becker, Amanda Z. Jiang, Dekker C. Deacon, David A. Nix, Melissa Q. Reeves, Kendell Clement, Robert L. Judson-Torres

## Abstract

Reliable chromosome segregation during mitosis is essential for high-fidelity mammalian cell division. Even human cancers burdened by whole genome duplication and aneuploidy generally retain sufficiently stable segregation to produce clonal populations of progeny. Cancer genomes are accordingly understood to evolve in punctuated bursts, in which catastrophic rearrangement is followed by clonal expansion of a stabilized karyotype^1–3^. However, bulk genomic sequencing, the workhorse of cancer genetics for the past two decades, cannot resolve structural variability within tumor subpopulations^3–5^. Here we identify a subset of cancers that exhibit continuous ultra-low fidelity chromosomal segregation, with single tumors harboring individual cells whose metaphase chromosome counts ranged from 25 to over 200. This remarkable phenotype is recurrent in multiple cancer types and most common in acral melanoma, where it is associated with dicentric chromosomes, translocations, and complex structural variants. Single-cell whole genome sequencing revealed ongoing structural variation and the genomic signature of microhomology-mediated break-induced replication (MMBIR), a backup DNA repair pathway that triggers chromoanasynthesis. These genomes therefore do not pass through isolated catastrophic events and settle into stable lineages but rather undergo continuous evolution of chromosome number and structure across generations, resulting in extreme intratumoral heterogeneity and a shifting repertoire of gene fusions. We anticipate these findings will serve as a starting point for classifying tumors by ongoing karyotypic and structural fluctuation and for testing whether this state confers selective dependence on backup repair pathways.

## Introduction

A tumor is usually described by its genome, and that description assumes there is one genome to describe. Bulk sequencing reads millions of cells at once and returns their average, which works if genomic events of interest are clonal or are present in large subpopulations of tumor cells. Even tumors that have duplicated their entire genome or lost whole chromosomes divide faithfully enough to pass a recognizable karyotype to their progeny, allowing detection of these chromosomal events. However, non-clonal chromosome changes, whole genome duplication, and wide ranges in chromosome number have remained difficult to resolve by whole exome or whole genome sequencing^2,4–6^. Complex rearrangements such as fold-back inversions are poorly captured by short reads, and structural events confined to a minority of cells are missed altogether^3–5^. Structural complexity in cancer is therefore modeled as something acquired early, maintained stably, and revised only in occasional punctuated bursts^1–3^. Recent work in model systems has challenged that model^7,8^, but it has rarely been tested directly in human tumors, because the tools to resolve rare structural variants have not been available.

Acral melanoma (AM) arises on palms, soles, and nail beds and is the predominant melanoma subtype in much of the world’s population. Unlike melanomas of sun-exposed skin, which respond to targeted and immune therapies, AM has limited effective and no tailored systemic treatments^9–11^. AM also carries one of the most unusual genomes described in solid cancer. It lacks the high ultraviolet-driven point mutation burden associated with other melanoma subtypes, and is instead defined by recurrent loci of extreme structural and copy-number complexity (‘SV/CNV loci’) containing *TERT*, *SKP2*, *RICTOR*, *CCND1*, *GAB2*, *YAP1*, *CDK4*, *MDM2*, and *CRKL* among other genes^12–18^. A single SV/CNV locus can be present in over 100 copies and participate in rearrangements that form a web of recursive fold-back inversions (FBI) interlaced with translocations to similarly affected loci on other chromosomes^12,18^. These SV/CNV loci have been called ‘tyfonas’^18^ and ‘hailstorms’^12^, which are descriptions that reflect their graphical visualization from bulk sequencing data. Yet despite these striking visualizations, the underlying rearrangements they represent have remained beyond the resolution of bulk approaches. AM shares these loci with a limited number of other tumors, including esophageal adenocarcinoma, dedifferentiated liposarcoma, and a subset of breast adenocarcinoma. How they arise is unknown^11,18^.

Given the prevalence of SV/CNV loci in AM and the comparatively late acquisition of single nucleotide variations^12,18^, we hypothesized that AM genomes evolve continuously through SV and CNV changes rather than stabilizing after an initial burst. Testing this has not been possible, because AM genetics has relied almost exclusively on short-read bulk tumor sequencing^12–18^. Here we combine a karyotype survey, reanalysis of existing AM genomes, and single-cell whole-genome sequencing (scWGS) to resolve SV and CNV evolution directly. Together they reveal karyotypes that drastically differ from cell to cell within single tumors, structural variants unique to every cell sequenced, and a rearrangement signature implicating ongoing microhomology-mediated break-induced replication, aka MMBIR (**Extended Data Figure 1**, visual abstract).

## Results

### Karyotypic fluctuation exceeds a whole genome in acral melanoma

Cytogenetic analysis of AM has been limited by the scarcity of models and the difficulty of establishing cell cultures^9,13^. Using models we previously derived, we generated metaphase spreads from six AM and three non-acral melanomas across eight patients, to our knowledge the largest such series reported (**Figures 1a-b, Extended Data Figure 2**, **Supplemental Data Table 1**, **Source Data 1**). Chromosome counts from cells within single AM models ranged from near-haploidy (<36 chromosomes) to hyperpentaploidy (>130 chromosomes), with HCI-AM085 containing cells that spanned 25 to over 200 chromosomes (**Figure 1b**, **Extended Data Figure 2a**). All AM karyograms contained neochromosomes, translocations, whole chromosome duplications and deletions, additions of unknown material to chromosome arms, and arm-level deletions. No extrachromosomal DNA (ecDNA) or double minute chromosomes were identified in any sample, excluding ecDNA amplification as a source of this variation.

**Figure 1:**
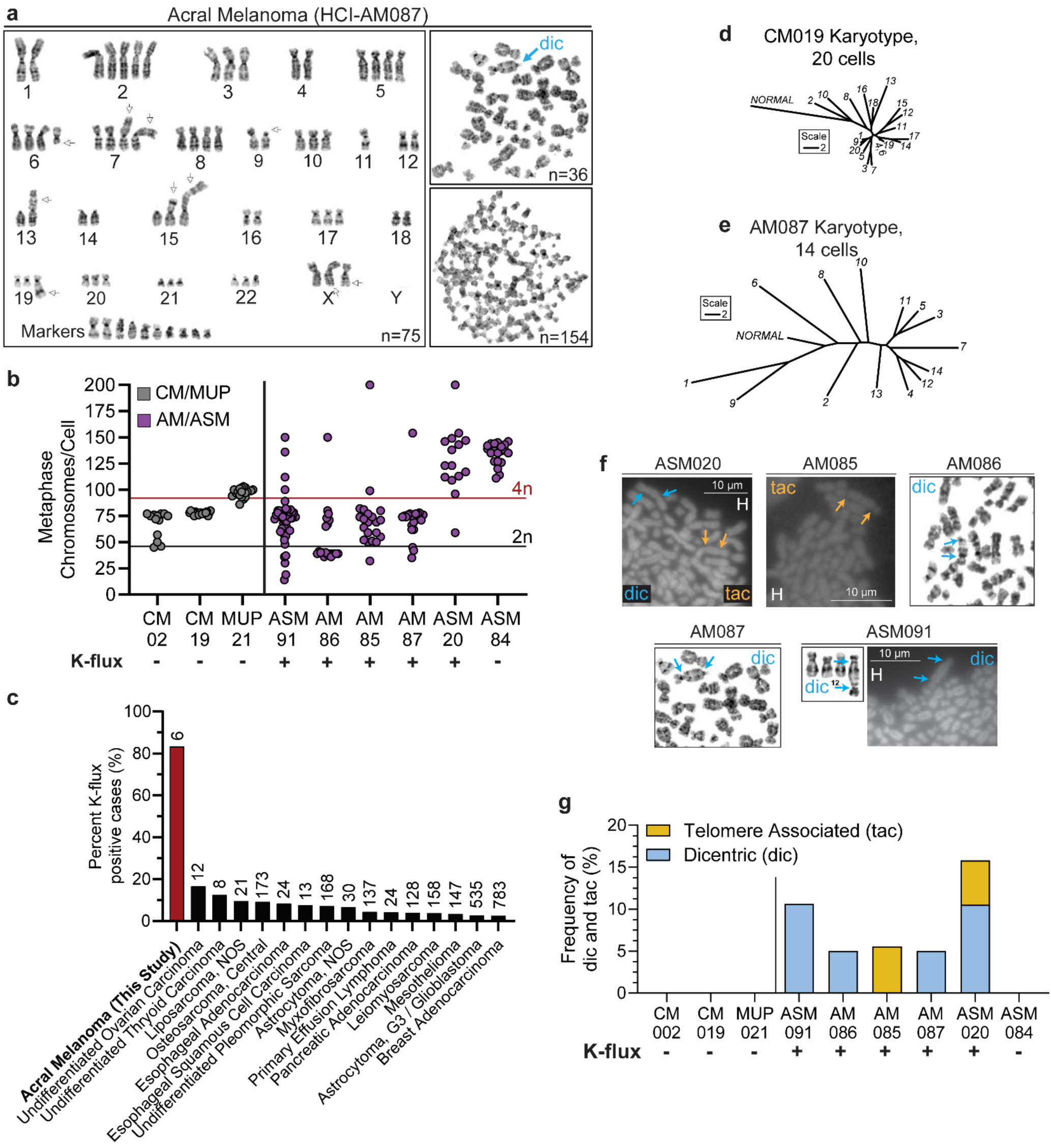
Individual cells within acral melanomas exhibit widely fluctuating chromosome numbers between cells (“K-flux”). **a**, Giemsa-stained AM karyotype and metaphase spreads for HCI-AM087 are shown. **b**, Chromosome counts from metaphase spreads in non-acral cutaneous melanoma (CM, gray), melanoma of unknown primary (MUP, gray), and AM cells (purple). Cells with >46 chromosome difference from maximum to minimum counts are labelled as K-flux positive under the x-axis. **c**, Cytogenetic cases from the Mitelman Database (n=84,506) were screened for K-flux positive status (black bars) and compared to our AM PDX cohort (red bar). The number above the bars represents the size of the cohort. **d**, Phylogenic trees show the degree of dissimilarity between individual karyotypes in CM model HCI-CM019, and **e**, AM model HCI-AM087. The scale measures increments of 2 different chromosome number or structure events and is uniform across all trees. **f-g**, Dicentric chromosomes (dic, cyan) and/or the non-covalent precursor telomere-associated chromosomes (tac, yellow) were enumerated from Giemsa or Hoescht (H) staining. Hoescht stained spreads have a 10 µm scale bar.

Five of six AM models showed differences in chromosome number count between cells exceeding 46 - the content of a normal 2n human genome. We term this striking intratumoral chromosomal variance ‘karyotypic fluctuation’, or K-flux (**Figure 1b**, **Extended Data Figure 2**, **Supplemental Data Table 1**).

The exception was the subungual model HCI-ASM084, which had undergone whole genome duplication and stably retained a >4n chromosome number, indicating that high ploidy alone does not produce K-flux (**Extended Data Figure 2h**). The non-acral controls, comprising a melanoma of unknown primary and cutaneous melanomas, were likewise K-flux negative (**Figure 1b**, **Extended Data Figure 2b-d**).

To assess K-flux frequency across tumor types, we screened the Mitelman Database of Chromosomal Aberrations and Gene Fusions in Cancer^19^ for cytogenetic reports where single tumors contained individual cells whose chromosome counts differed by more than 46 chromosomes from one another (n=84,506). K-flux positive cases were rare overall but recurred in a subset of malignancies (**Figure 1c**). Notably, several of the cancer types where this was most common, such as breast adenocarcinoma, esophageal adenocarcinoma, and dedifferentiated liposarcoma, have been reported to harbor SV/CNV loci^18,20^. Most of these malignancies, including esophageal adenocarcinoma, undifferentiated pleomorphic sarcoma, osteosarcoma, dedifferentiated liposarcoma, mesothelioma, and primary effusion lymphoma, share similarly poor survival and therapy response^21^. AM was the most frequently K-flux positive of any cancer type examined.

K-flux describes variance in chromosome number but does not factor for other differences in chromosome structure. We therefore built phylogenetic trees from individual karyograms of each model, using chromosome number together with marker chromosomes, translocations, additions, deletions, derivatives, isochromosomes, breaks, and dicentrics (**Figures 1d-e**, **Extended Data Figure 2i-j**). Spreads from the non-acral models clustered on short branches, consistent with stable dominant clones. Spreads from HCI-ASM084 and HCI-AM087 were separated by long branches, indicating that individual cells had diverged substantially. In HCI-ASM084 (**Extended Data Figure 3j**), this divergence arose largely from random small gains and losses of whole chromosomes, the expected product of ongoing chromosomal instability. In HCI-AM087 (**Figure 1e**), it arose instead from cell-specific chromosome additions and varying numbers of neochromosomes, with the hypodiploid and near-haploid cells segregating from the remaining near-diploid and near-triploid population. The largest HCI-AM087 spread (154 chromosomes, **Extended Data Figure 2a**) was too low resolution to include, so this tree understates the variation present.

We next asked whether the karyotypic variation seen in culture was also present in tumor tissue. Nuclear ploidy was measured by propidium iodide flow cytometry of nuclei extracted from FFPE sections of patient-derived xenograft tumors. The three CM tumors had G0/G1 dominant profiles matching a normal diploid control. Of nine AM tumors, only HCI-ASM020 shared this pattern, while the remaining eight showed abnormal profiles with hypertetraploid populations, a dominant G2/M or tetraploid peak, or both (**Extended Data Figure 3**). This variation in tumor tissue is consistent with that seen in transient and long-term culture. Fluctuation of this magnitude requires a continuing source of chromosome breakage, which led us to examine telomere integrity.

### Karyotypic fluctuation occurs only in AM models with chromosome fusions

The karyograms and metaphase spreads revealed recurrent dicentric chromosomes in AM (‘dic’ in **Figures 1a and 1f**, **Extended Data Figure 2g**, **Supplemental Data Table 1**, **Source Data 1**). All K-flux positive AM models carried dicentrics or telomeric associations - the non-covalent precursor structure in which joined chromosomes remain separated by a visible gap^22,23^. Both structures form as telomeres shorten (**Extended Data Figure 4a**). No K-flux negative CM or AM model carried either (**Figure 1g**, Fisher’s exact p=0.0008).

The linkage of two active centromeres, as found in dicentric chromosomes, is mechanically incompatible with normal segregation. During anaphase the spindle pulls the joined chromosomes toward opposite poles, producing an anaphase bridge that stalls mitosis and can initiate breakage-fusion-bridge (BFB) cycles^24–26^. Consistent with this, anaphase bridges quantified by DAPI staining were more frequent in AM than in CM or non-cancer controls (one-sided exact permutation Mann-Whitney, p=0.0429) (**Extended Data Figure 4b-c**). AM models showed no increase in micronuclei or nuclear buds relative to normal or CM controls (**Extended Data Figure 4d**), consistent with bridge resolution occurring in or adjacent to the primary nucleus, as described during telomere crisis^27,28^, rather than through the micronucleus-mediated route to chromothripsis^26,29^.

Dicentrics were present in 16% of HCI-ASM020 spreads and this model was K-flux positive despite it retaining a diploid DNA content profile by flow cytometry (**Extended Data Figure 3b**). This indicates that dicentric formation can alter chromosome number in individual cells without shifting the average ploidy of the population. Together, the persistence of chromosome fusion events and their disruption of mitotic segregation support a model in which fusion is tightly linked to K-flux in AM.

### SV/CNV loci are unstable in AM clinical cases

As chromosome number and structure are unstable within AM cells, we asked whether copy number changes at SV/CNV loci reflect gain or loss of whole chromosome arms or the generation of new structural events within the loci themselves. Copy number was tracked across genes recurrently present within SV/CNV loci, including *CDK4, CRKL, GAB2, MDM2* and *TERT*, in pairs of AM tumors collected from the same patients (**Extended Data Figure 5a**)^12–16,18^. During the several months between collection of HCI-AM086 and HCI-AM087, the *CDK4* and *GAB2* loci underwent further amplification (Mann-Whitney U, p<0.0001) (**Figure 2a**). Between the HCI-AM090 primary tumor and its associated HCI-AM135 lymph node metastasis, *CRKL* and *TERT* instead lost copies (p=0.0002 and p<0.0001) (**Extended Data Figure 5b**). The same instability was apparent when average copy number at SV/CNV loci was followed in long-term culture (**Extended Data Figure 5c-d**).

**Figure 2:**
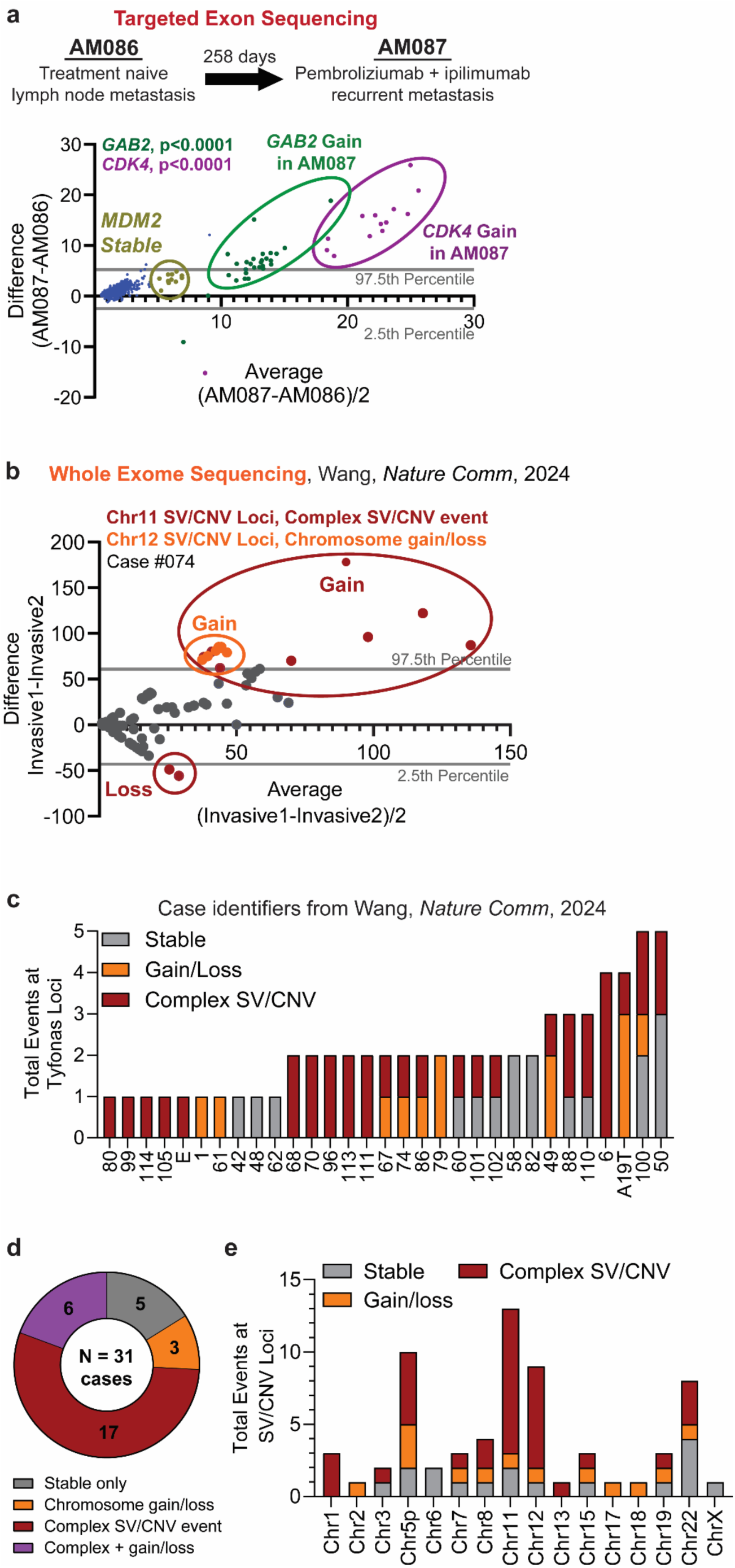
Clinical AM specimens demonstrate dynamic SV/CNV. **a**, HCI-AM086 and HCI-AM087 were collected from the same patient before and after treatment. Copy number from targeted exon sequencing is compared using a Bland-Altman plot. The most amplified genes, *GAB2* and *CDK4*, underwent an increase in copy number in HCI-AM087 (p<0.0001, Mann-Whitney U). **b**, The same analysis was performed on copy number segments from a study with serial whole exome sequencing (WES) of clinical AM cases^12^. In exemplar case #074, the Chr12 SV/CNV loci had whole chromosome segment gain/loss (all segment outliers present outside one confidence interval), and the Chr11 SV/CNV loci developed a new complex SV/CNV loci event (segments spread outside of both confidence intervals). All cases are shown in Extended Date Figure 5. **c-d**, The majority of clinical cases had ongoing complex SV/CNV events, which **(e)** were enriched at Chr1, Chr5p, Chr11, Chr12, and Chr22.

We performed a similar analysis of WES from AM specimens serially acquired from the same patients^12,14^. Two patterns were distinguishable. Concordant shifts in copy number across a chromosome arm are consistent with gain or loss of that arm and imply no restructuring within the loci. Opposing shifts between loci on the same chromosome cannot be explained by whole-chromosome dosage and instead imply a new structural or complex copy number event, which we term a complex SV/CNV event. Case 074 of Wang et al^12^ displays both: all chromosomal segments associated with the Chr12 loci were uniformly amplified between Invasive1 and Invasive2, consistent with chromosome gain, whereas segments from the Chr11 loci were found shifted in opposing directions (**Figure 2b**). Across the full cohort, 23 of 31 cases (74%) carried a complex SV/CNV event at one or more loci, while only 5 of 31 (16%) were stable and 3 of 31 (10%) showed arm-level gain or loss alone (**Figure 2c-d**, **Extended Data Figure 5e-ak**). The chromosomes undergoing complex SV/CNV evolution were also those carrying the most SV/CNV loci, namely Chr5p, Chr11, Chr12 and Chr22 (**Figure 2e**). Compared with the remainder of the exome, SV/CNV loci changed significantly in 79% of paired cases and in 73% of pairwise comparisons among cases with three tumors (**Extended Data Figure 5aj-ak**).

A second published cohort gave the same qualitative result, with 4 of 11 paired cases showing complex SV/CNV evolution between paired tumors and another 4 showing arm-level change (**Extended Data Figure 5ai-ax**)^14^. The lower rate relative to the first cohort likely reflects that these data lacked SV/CNV loci annotation, requiring us to call loci from amplification thresholds and reducing sensitivity. In both cohorts, SV/CNV loci evolve during disease progression rather than remaining fixed once formed.

### SV/CNV loci are formed from chromoanasynthesis (CAS) events

The etiology of SV/CNV loci in AM has been debated since their discovery^15,18^, though most reports attribute them to a complex and catastrophic chromoanagenesis (Greek: ‘chromosome rebirth’) event^12,18^. Chromoanagenesis types includes chromothripsis (‘chromosome shattering’), chromoplexy (‘chromosome restructuring’), and chromoanasynthesis (CAS or ‘chromosome reconstitution’)^30^, which is driven by distinct underlying molecular mechanisms. To determine which type generated these loci in AM, we conducted WGS of HCI-AM087 and identified SV/CNV loci using JaBbA^18^. Consistent with a prior report^18^, we found FBI and translocations involving multiple loci across the AM genome (red and purple lines in ‘tyfonas’ pattern in **Extended Data Figure 6**). The observed amplified segments remained in chromosomal context, as neither extrachromosomal nor circular amplicons were identified, consistent with the absence of double minutes from metaphase spreads. Copy number across the Chr11 and Chr12 SV/CNV loci rose progressively to 40 copies through many intermediate states. This pattern is distinct from that expected from chromothripsis, in which shattered fragments are randomly re-ligated and yield a copy number profile that oscillates between neutral copy number or various states of deletion. The loci were also strongly unbalanced, with large net copy gain. This is also inconsistent with chromoplexy which produces translocations and smaller, rare deletions from re-ligation without amplification. Thus the observed progressive copy gain, built from recursive local FBI and interchromosomal translocations, best fit the expected chromosomal architecture of CAS^30–33^.

CAS occurs when stalled DNA replication forks or DNA breaks are repaired by MMBIR. This form of DNA repair relies on DNA template switching to bypass the damage and restart DNA synthesis. Polymerases anneal to regions of 0-10 bp of microhomology in spatial proximity to the stalled fork, copy a short distance, and then anneal to a new tract of microhomology, repeating until normal DNA synthesis can resume^30–33^. Because the mechanism copies rather than ligates, multiple template switches produce recurrent local FBI leading to high SV and CNV, as well as translocations between chromosomes where microhomology is shared. MMBIR therefore predicts three features at AM SV/CNV loci: microhomology at breakpoint junctions, FBI separated by characteristic short distances, and translocations seeded by homology at distal sites.

To assess these features, scWGS was performed on 14 freshly harvested HCI-AM087 cells as well as five HCI-CM019 as controls using primary template-directed amplification on individual viable cells from PDX tumors (**Figure 3a**, **Extended Data Figure 7**). We assessed intrachromosomal FBI in *GAB2*, one of the most frequently involved genes in SV/CNV loci^12,16^ and highly amplified in HCI-AM087 (**Extended Data Figure 8a-c**). An example FBI from HCI-AM087 cell 10 is shown in **Figure 3b**. In this example, three serial breakpoint junctions with 4-10 bp of microhomology overlap were identified within a 600 bp region. Across all AM cells there was an increased number of FBI within *GAB2* compared to HCI-CM019 (**Figure 3c**, Mann-Whitney U, p<0.0001, **Supplementary Data Table 2**). The majority of FBI junctions had breakpoint mates located <100 bp or between 300 and 450 bp away (**Figure 3d**), aligning with reports that most MMBIR-induced breakpoint pairs in normal human fibroblasts and other model organisms are separated by 10-700 bp^32–35^. Of these intrachromosomal FBI, 13% of the alternative sequences at breakpoints also had high or perfect homology to regions on at least one other chromosome, providing the microhomology needed for interchromosomal translocation template switching during MMBIR. Strikingly, 95.7% of FBI occurred between intronic sequences, rendering them invisible to WES.

**Figure 3:**
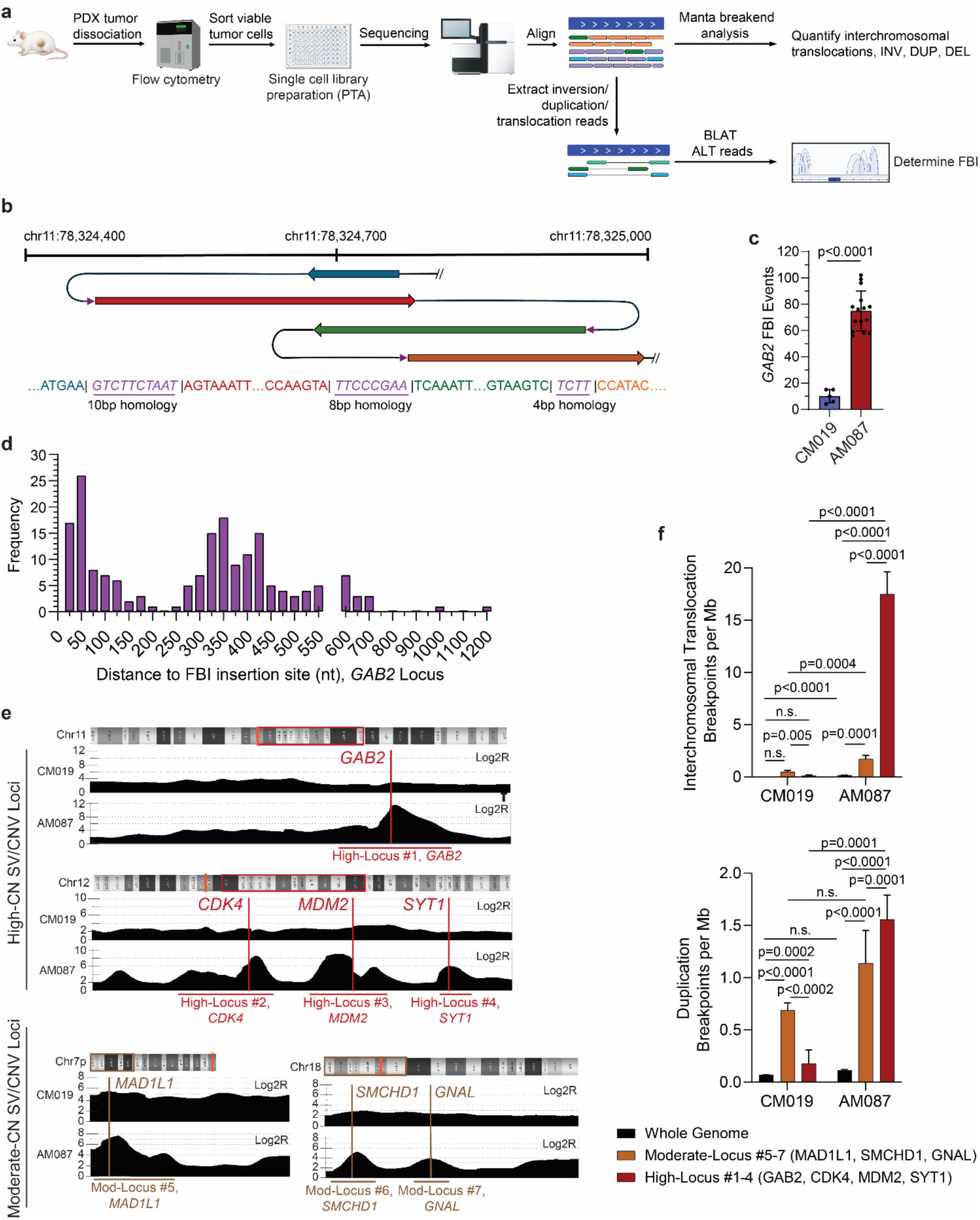
High-CN SV/CNV loci (High-Loci) loci carry the microhomology fold-back inversion signature of chromoanasynthesis (CAS). **a**, HCI-AM087 and HCI-CM019 PDX tumors were harvested, depleted of mouse stroma, and sorted as single viable cells for primary template-directed template amplification (PTA) library preparation on 5x CM019 and 15x AM087 cells. **b**, FBI within the *GAB2* locus were manually interrogated with BLAT on breakpoint ALT-read residues to identify the connecting intrachromosomal breakend location. A representative set of two FBI from an AM cell (AM-c10) are shown with the bridging microhomology written in purple text. **c**, HCI-AM087 cells have an increased frequency of unique FBI events within *GAB2* compared to HCI-CM019 (Mann-Whitney U, p<0.0001). Each dot represents a single cell genome. **d**, These intrachromosomal FBI favor short distance references with bimodal peaks at <100 and 300-450 nt away from the breakpoint, consistent with CAS scarring^32,34^. **e**, OncoScan CNV array copy number plots for High-Loci (red) and Moderate-CN SV/CNV loci (Mod-Loci or Moderate-Loci, brown) are shown. High-Loci can be identified by CNV array alone, and Moderate-Loci require WGS to resolve. The full chromosome banding pattern is shown above each plot with centromeres represented by orange bands and the plotted regions by the red or brown boxes. The location of representative genes for each locus is shown by the vertical colored lines. **f**, Comparing AM to CM, the dominant SV/CNV loci breakpoints profiles are the interchromosomal translocations and duplications (normal distribution by Shapiro-Wilk test, paired t tests with a minimal significance cutoff of p<0.001).

We next compared breakpoint types across 10 genomic loci, ranging from 4.5 to 10 Mb in size, showing varying degrees of SV/CNV by copy number (CN): four high-CN SV/CNV loci on Chr11 and Chr12 containing *GAB2*, *MDM2* and *CDK4* (High-Loci); three moderate-CN loci on Chr7p and Chr18 identified as SV/CNV loci only by WGS (Moderate-Loci); and three simple loci on Chr1p, Chr4p and Chr8q not classified as SV/CNV loci by WGS (Simple Loci) (**Figure 3e, Extended Data Figure 8c-h**). Translocations and tandem duplications were significantly enriched in both the High- and Moderate-Loci relative to the whole genome and to Simple Loci (**Figure 3f, Extended Data Figure 9a-b**), and High-Loci showed a 7-fold increase in both translocations and tandem duplications compared with the same regions in CM (**Figure 3f**). Insertions and deletions were not significantly different between CM and AM at these loci (**Extended Data Figure 9c-d**). This enrichment for translocation and tandem duplication breakpoints without accompanying deletion or insertion enrichment is characteristic of CAS, and reinforces the architectural evidence against chromothripsis and chromoplexy, both of which generate deletion- and insertion-rich junction profiles. Together, these data indicate that SV/CNV loci in AM are products of CAS driven by MMBIR.

### SV/CNV loci fluctuate between individual cells

If SV/CNV loci are products of CAS, we asked whether that process had completed or, like K-flux, remained ongoing. Comparing breakpoints across the same 10 loci, the number of interchromosomal translocations varied significantly between individual AM cells, but only in the High-Loci (**Figure 4a**, two-way ANOVA p=0.001, **Extended Data Figure 9e-i**). Moderate- and Simple Loci in AM, and all loci in CM, were stable. The translocation partners were equally variable. Translocations originating in the High-Loci connected to most chromosomes, with the densest connections to Chr7, 11, 12 and 19, and no two AM cells shared the same pattern (**Figure 5b**, **Extended Data Figure 9j-s**, **Source Data 2**). Ultimately, every AM cell possessed a distinct interchromosomal translocation profile.

**Figure 4:**
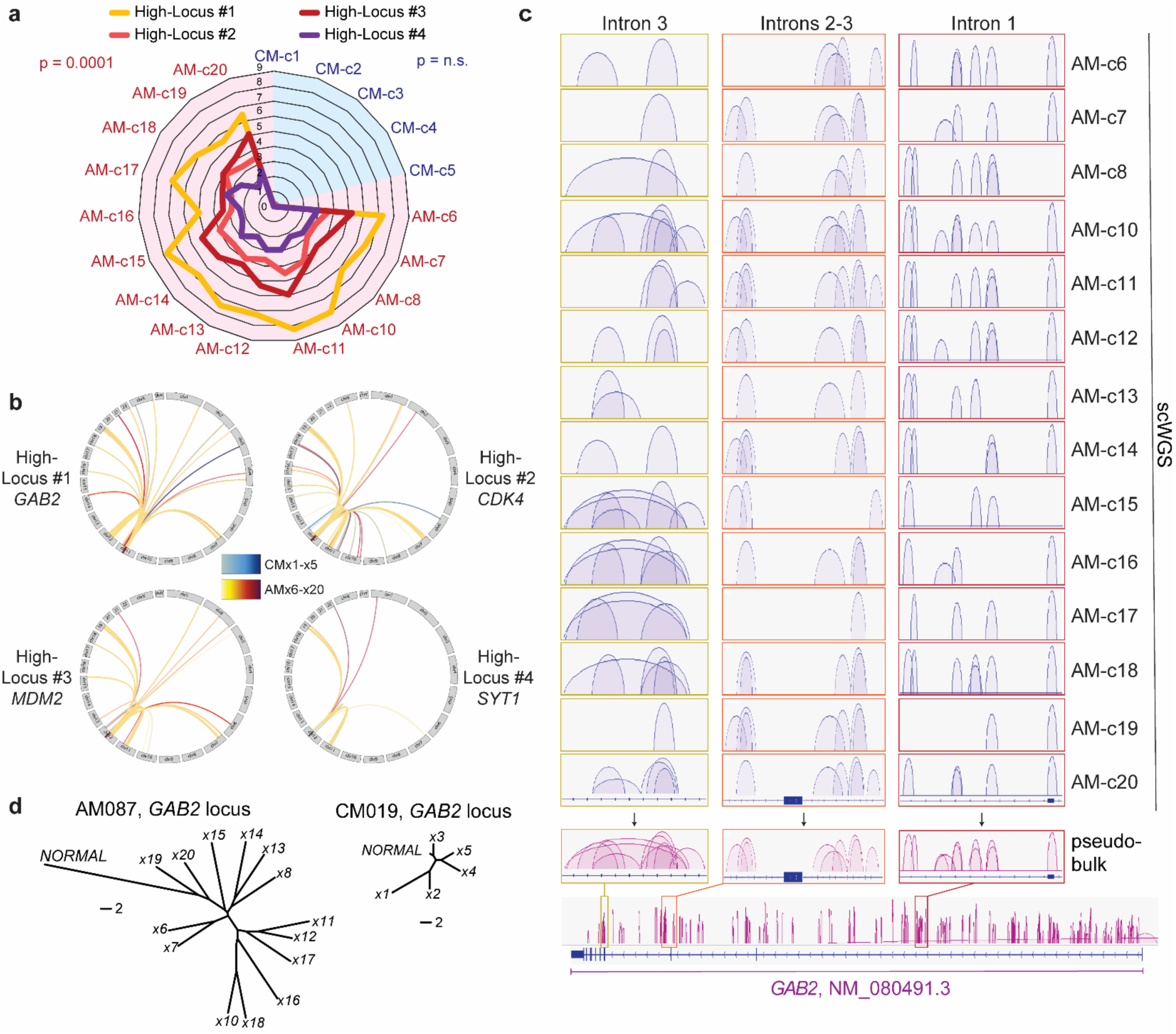
Individual AM cells have unique translocation and FBI fingerprints, indicating ongoing chromoanasynthesis (“CAS-flux”). **a**, The frequency of translocation events at High-Loci loci varies significantly between AM cells across High-Loci (2-way ANOVA, p=0.0001). Each ring radiating from the center represents the number of translocations per Mb. HCI-CM019 are color coded blue, and HCI-AM087 are red **b**, Aggregate circos plots of all single cell genomes for each locus summarize the translocation heterogeneity between cells. HCI-AM087 cells are warm colors (yellow, orange, and red), and HCI-CM019 cells are cold colors (blue, grey, and purple). Individual circos plots per cell and locus are presented in **Extended Data Figure 9j-s**. **c**, Each AM cell had a different FBI fingerprint (blue sashimi plots) and, when amalgamated (magenta sashimi plot), revealed a predilection for intronic sequences. The exon sequence in intron 1 is an alternative transcript start site. **d**, Phylogenetic plotting of these differences reveal that individual AM cells have a greater variation in FBI fingerprints compared to CM cells.

**Figure 5:**
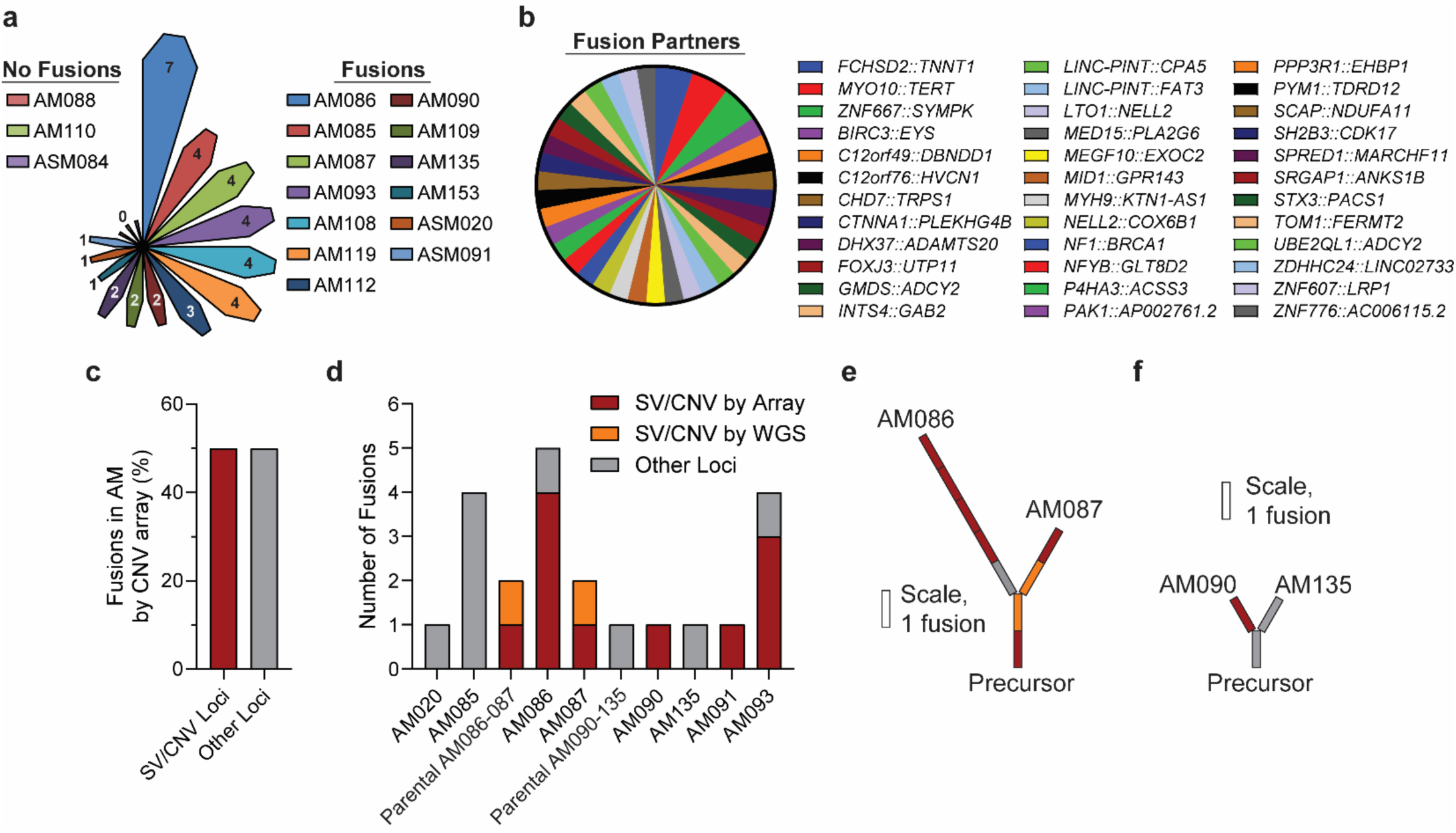
Gene fusion events involve SV/CNV loci, explaining the prevalence and variety of gene fusions in AM. **a**, The majority of AM have at least 1 gene fusion as identified by PDX tumor RNA-seq. Two of the fusionless models, HCI-AM088 and HCI-ASM084, have no suspicious regions for SV/CNV loci by CNV array. **b**, Most fusion partners are non-cancer associated genes. A minority comprises oncogenes such as *GAB2* and *TERT* or tumor suppressors (*NF1*, *BRCA1*). **c**, Half of gene fusions occur within SV/CNV loci, as identified by CNV array. **d**, The number and type of locus involved by fusions events are shown according to model. Conserved fusions within patient-matched models are separated from fusions occurring in only one model. For HCI-AM087, fusions at orange SV/CNV loci were only identified by WGS. **e-f**, Patient-matched PDX models demonstrate evolution of gene fusions from a common precursor, consistent with the CAS-flux model.

We then asked whether an independent class of rearrangement showed the same instability. The FBI events characterized in *GAB2* were visualized per cell and as a pseudobulk composite (**Figure 4c**, **Supplemental Data Table 2**). No two cells carried the same FBI fingerprint, and HCI-AM087 showed a far wider phylogenetic spread of FBI number and position than HCI-CM019 (**Figure 4d**). Two independent rearrangement classes, translocations and FBIs, therefore differ between every AM cell examined. Because no ecDNA or double minutes were detected in any model (**Figure 1**, **Extended Data Figure 2**, **Source Data 1**), this diversity resides in linear chromosomal loci rather than in independently segregating extrachromosomal elements. Together with the CN changes between paired patient tumors (**Figure 2**), these data show that SV/CNV loci are not merely complex but continue to fluctuate in number and position, a process we term CAS fluctuation, or CAS-flux.

### SV/CNV loci are detectable by a clinical copy number array

SV/CNV loci are currently defined by WGS, which is not routinely available in clinical practice. To assess whether they can be identified by a deployed assay, we mapped the four High-Loci called by JaBbA in HCI-AM087 onto the OncoScan CNV array. These loci produce a distinctive asymmetric peak and plateau on the log2R copy number track (**Extended Data Figure 8**, **Source Data 3**), from which we derived array-based identification criteria (see **Methods**). These criteria identified SV/CNV loci in most remaining models (**Extended Data Figures 8g-h**), converging on the same recurrently amplified regions identified by targeted exon sequencing in Chr11 (*GAB2*), Chr12 (*CDK4*, *MDM2*) and Chr22 (*CRKL*). HCI-ASM084, the only K-flux negative model, was also the only model in which no SV/CNV loci could be identified, a concordance consistent with a shared origin for the two phenomena. The array was limited to high-CN loci: regions of moderate-CN loci such as Chr7p, Chr16p and Chr18p in HCI-AM087 could not be distinguished from simple amplification events without WGS (**Extended Data Figures 8c-d**). Clinical CNV array therefore offers a tractable route to detecting the most prominent high-CN SV/CNV loci.

### CAS-flux generates AM gene fusions

Acral melanomas are enriched for gene fusions^15,36^, but the generative mechanism has not been identified. Because a translocation that joins two transcribed genes produces a fusion transcript, we hypothesized that SV/CNV loci in AM should be enriched for fusions, and fusion accumulation should be on-going. We performed RNA-sequencing on 16 AM, 48 CM, 18 MUP and 3 mucosal melanoma PDX models to identify gene fusion events (**Figure 5**, **Extended Data Figure 10a-c**, **Supplementary Data Tables 3-4**, **Source Data 4**). Most AM tumors carried at least one fusion (81%, 13/16), with HCI-AM086 carrying seven (**Figure 5a, Extended Data Figure 10d-e**). None were recurrent **(Figure 6b**). *MYO1::TERT* was shared by HCI-AM090 and HCI-AM135, and *FCHSD::TNNT1* and *ZNF66::SYMPK* by HCI-AM086 and HCI-AM087, each pairs of tumors from the same patient and consistent with inheritance from a common ancestor rather than recurrent independent formation. Only 36% of fusions involved genes with known or suspected roles in oncogenesis (**Supplemental Data Table 4**), although at least one such fusion was present in 12 of 16 tumors. The predominance of passenger fusions indicates that fusions in AM arise frequently rather than rarely, since a rare process would yield mostly events retained by selection.

**Figure 6:**
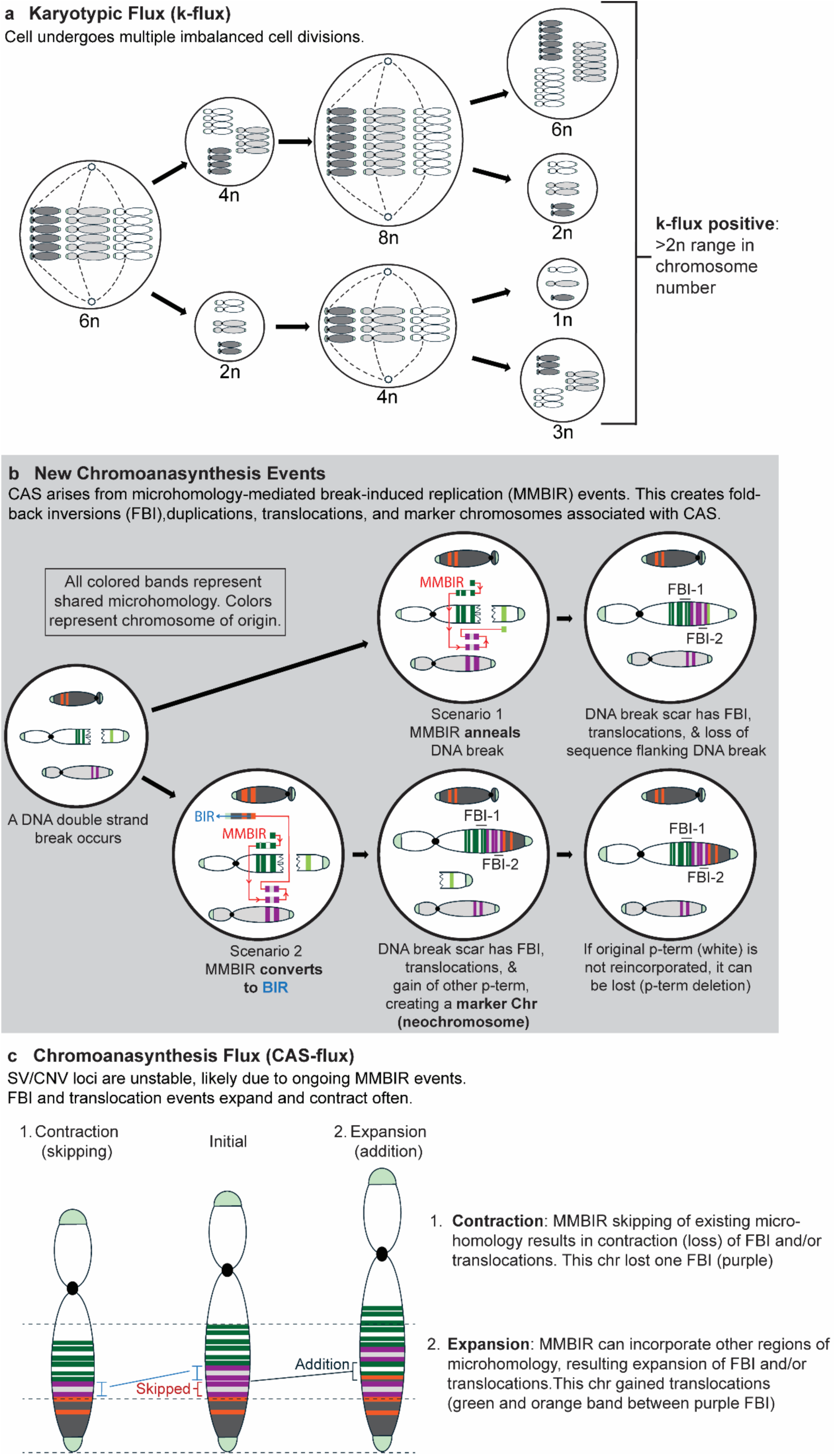
CAS-flux and K-flux drive perpetual genomic change and extreme intratumoral heterogeneity. **a**, A 6n cell from a K-flux positive tumor undergoes imbalanced cell divisions. After two divisions, there is spread of cells spanning a 5n range of chromosome counts. The presence of dicentric or telomere-associated chromosomes may trigger this chromosome nondisjunction (not shown). Small black circles are centrioles, dashed lines are microtubules, and light green chromosome ends are telomeres. **b**, CAS is a complex SV/CNV event that occurs when DNA is repaired with MMBIR. Low processivity DNA polymerases perform template switching at regions of micohomology (bands that are colored according to chromosome of origin) to bridge the double strand break. In both examples, MMBIR first template switches to an intrachromosomal microhomology, forming an FBI (green). It then template switches to a microhomology on a different chromosome (purple), forming a translocation. Another template switch creates a second FBI (purple). In scenario 1, MMBIR successfully anneals the DNA break by template switching to a microhomology on the terminal white chromosome fragment. All intervening sequence between the initial green microhomology and the lime green microhomology is lost. In scenario 2, MMBIR fails to bridge the break and instead converts to BIR after track switching to the orange microhomology. DNA synthesis copies the remainder of the other chromosome, leading to an amplification of this region and the generation of a marker chromosome (aka neochromosome). If the terminal white fragment is not reincorporated into a chromosome, it can be lost (p-terminal deletion). **c**, FBI and translocation events within SV/CNV loci evolve with each cell division, likely due to ongoing MMBIR repair. This can lead to 1) contraction of the number of FBI if MMBIR skips over existing intrachromosomal microhomology. Alternatively, 2) expansion occurs if MMBIR template switches to other intra- or extrachromosomal regions of microhomology.

Fusion status tracked with the presence of SV/CNV loci. Among the nine AM models with both CNV array and RNA fusion data, all seven models with SV/CNV loci also carried fusions, and the two without SV/CNV loci (HCI-AM088 and HCI-ASM084) did not (9/9 concordant, Fisher’s exact p=0.028). HCI-ASM084, the one K-flux negative model of the six karyotyped, is therefore negative for all three features, while HCI-AM088 failed to grow in culture and could not karyotyped (**Supplementary Data Table 5**). Half of all AM fusions fell within confirmed SV/CNV loci, a greater than 14-fold enrichment over the 3.4% of the genome these loci occupy (exact binomial p=2x10⁻¹¹) (**Figure 5c**). This is a conservative estimate: in HCI-AM087, where WGS was available, two of four fusions at SV/CNV loci would have been scored as simple amplifications by array alone (**Figure 5d**).

Where multiple tumors were available from the same patient, fusions were discordant, with private fusions acquired in post-treatment samples and in metastatic disease (**Figures 6e-f**). In HCI-AM086 and HCI-AM087, private fusions involving the cancer-associated genes *NELL2*, *PAK1* and *ZNF776* were acquired only after the branch point. In these models and HCI-AM090 most new fusions fell within SV/CNV loci. Fusions in AM are therefore not a fixed feature of the genome but continue to arise at these loci as disease progresses, the expected signature of CAS-flux rather than of a completed event.

## DISCUSSION

We describe two forms of ongoing chromosomal evolution in acral melanoma (**Figure 6**, **Extended Data Figure 1**). K-flux is a repeating series of non-disjunction events producing a spread of more than a whole genome in chromosome number across a cell population. CAS-flux is the continuous restructuring of SV/CNV loci through MMBIR. Neither is visible in bulk tumor sequencing, and each was detectable only below the bulk average: in single cells, metaphase spreads, or matched-pair comparison of amplified loci.

Why CAS-flux persists rather than resolving might be explained by the structures it creates. MMBIR resolves stalled or collapsed replication forks by template switching at microhomology, and in doing so generates FBI. Because of their repetitive nature, FBI are predisposed to forming DNA secondary structures that themselves stall replication forks^30,37,38^. We therefore propose that SV/CNV loci are a positive feedback loop: MMBIR resolves replication problems at secondary structures, introduces additional FBI and secondary structures, and increases reliance on MMBIR at the same loci. This accounts for the variable SV breakpoints between individual cells (**Figure 4**), the continued acquisition of gene fusions (**Figure 5**), and the concentration of both at a small number of recurrent loci.

The relationship between K-flux and CAS-flux is unresolved. They co-occur in every model examined, and HCI-ASM084 was negative for both, which is consistent with a shared origin. Dicentric chromosomes offer one candidate link since their missegregation generates both aneuploidy and the double-strand breaks that MMBIR repairs. However, this connection has been difficult to establish in human models^39^. Aneuploidy and whole genome duplication alone are insufficient to produce non-disjunction of this magnitude^40,41^, so an upstream process remains to be identified.

Our findings do not contradict the observation that SV/CNV loci are clonally shared across tumors from the same patient^12^. Our data agrees that they are shared, and once an unstable SV/CNV locus is formed, the tumor will never reestablish a normal chromosomal segment in that locus. Matched-pair WES, karyotyping, and scWGS additionally allowed us to ask whether the exact states of those loci shift over time, and they do. Specific copy number states, translocations, FBI, and fusion partners continue to evolve amongst the cell population without destroying the SV/CNV loci. This phenomenon is potentially a more common oncogenic process, and our findings argue that further study of it is needed. Dedifferentiated liposarcoma, esophageal adenocarcinoma and a subset of breast adenocarcinoma harbor SV/CNV loci^18^ and appear among the K-flux positive types in cytogenetic records, making them the first places to ask whether CAS-flux similarly co-occurs.

CAS-flux also accounts for two unexplained features of AM genetics. First, the non-APOBEC hypermutation that is concentrated within SV/CNV loci^15,18^ is expected if those loci are repeatedly copied by the low-fidelity MMBIR polymerases, such as Polζ and Polθ^32,42^. Second, marker chromosomes, which we observed in every AM karyogram examined (**Figure 1**, **Extended Data Figures 2**), are the expected product when a template switch to another chromosome converts to BIR^37,42^.

CAS-flux places AM genomes in continuous reliance on MMBIR, a pathway that normal cells use only to rescue unresolved DNA lesions prior to mitosis. This nominates the low-fidelity polymerases that carry out MMBIR as a candidate vulnerability. Prior work raised the possibility that sustained instability in AM would implicate an exploitable repair defect but did not pursue it, having concluded the instability was punctuated and self-limiting^12^. Inhibition of the low-fidelity polymerases that carry out MMBIR is the direct test of that hypothesis.

Two conventions have previously obscured these processes. Clinical karyotyping focuses on identifying clonally related populations, which suppresses reporting of extreme near-haploid or hypertetraploid karyograms (ISCN 2024, section 4.5.11), so the K-flux frequencies recorded in cytogenetic catalogues are likely underestimates. Similarly, bulk sequencing averages across millions of cells, so structural variation confined to subpopulations is absent from the record by regression to the mean. A genome described as structurally stable may simply have been measured at a resolution that cannot detect active remodeling. The combination of single-cell sequencing, cytogenetics, and matched-pair analysis applied here provides a way to ask how widely that is true.

## METHODS

See Supplemental Information section.

## ACKNOWLEDGEMENTS

We thank Leo Lin and Elizabeth Anderssen from ARUP Laboratories; Evan Martin and Elizabeth Gonzalez from Karyologic Inc; and Katie Kennedy and John Bon from Bioskryb Genomics. Core resources used include ARUP Clinical Trials Office, HCI High Throughput Genomics, HCI Cancer Bioinformatics Shared Resource, HCI Preclinical Model resources, HCI Immuno-Onocology Network, HCI Total Cancer Care, supported by the National Cancer Institute of the National Institutes of Health under Award Number P30CA042014. The computational resources used were partially funded by the NIH Shared Instrumentation Grant 1S10OD021644-01A1. The content is solely the responsibility of the authors and does not necessarily represent the official views of the University of Utah or the funding agencies.

## AUTHOR CONTRIBUTIONS

Conceptualization: EAS

Data curation: EAS, AP, JZ

Formal analysis: EAS, AP, RGZ, MH, JZ, MQR

Funding acquisition: EAS, RLJT

Investigation: EAS, AP, RGZ, CAB, MH, AZJ

Methodology: EAS, AP, JZ, CAB, MQR, DN, KC

Project administration: EAS, DCD, RLJT

Resources: EAS, AP, RGZ, CAB, AZJ

Software: MQR, DN, KC

Supervision: EAS, DCD, RLJT

Validation: EAS, AP, JZ, DN, KC

Visualization: EAS, AP, JZ, MQR, DN, KC

Writing – original draft: EAS, MQR, RLJT

Writing – review and editing: all authors

## MATERIALS AND CORRESPONDENCE

All requests for materials and correspondence should be made to Dr. Robert Judson-Torres.

## Supplementary Information

### Methods

#### PDX tumor growth and dissociation

Patients donated tumor tissue after written consent as part of the University of Utah IRB 89989 and Huntsman Cancer Institute (HCI) IRB 10924. PDX tumors were maintained, propagated, and characterized as described previously^13^. A list of all PDX models used in this study is tabulated in **Supplementary Data Table 5**. Tumor cell dissociation was performed by the Huntsman Cancer Center (HCI) Immuno-Oncology Network resource using Milteny dissociation enzyme mix (130-095-929) on a gentleMACS™ Octo Dissociator (Milteny, 130-096-427) with the 37C_h_TDK_1 protocol. Mouse cell depletion was performed using the mouse cell depletion kit buffer and beads (Milteny, 130-104-694) on an autoMACS® Pro Separator (Milteny, 130-092-545) equipped with an autoMACS column (Milteny, 130-021-101) using the DEPLETES program. Full protocol details have been published previously^13^.

#### Cell culture

A full list of cell culture models is in **Supplementary Data Table 5**. All cells were cultured at 37°C and 5% CO_2_. Primary human melanocytes were cultured in Medium 254 (Gibco, M254500) supplemented with Human Melanocyte Growth Supplement (Gibco, S0025). HCI-AM087, HCI-ASM091, HCI-CM002, and HCI-CM019 were cultured in Mel2 medium, comprised of 400ml MCDB 153 Medium (Sigma-Aldrich, M7403-10X1L), 100ml Leibovitz’s L-15 Medium (Gibco, 11415-064), 2.5% fetal bovine serum, 1.68mM CaCl2, 5ml Insulin-Transferrin-Selenium-Ethanolamine (ITS-X) (Gibco, 51500-056), 5ng/ml Epidermal growth factor (Sigma-Aldrich, E-9644), 15µg/ml bovine pituitary extract (Gibco, 13028-014), and 1x Penicillin-Streptomycin (Gibco, 15140122). All other AM cells were cultured in AM3 medium, comprised of 80% MCDB 153 (Sigma-Aldrich, M7403-10X1L), 20% Leibovitz’s L-15 Medium (Gibco, 11415-064), 2.5% fetal bovine serum, 1.68mM CaCl_2_, 1x Insulin-Transferrin-Selenium-Ethanolamine (ITS-X) (Gibco, 51500-056), 5ng/ml Epidermal growth factor (Sigma-Aldrich, E-9644), 0.2% bovine pituitary extract (Gibco, 13028-014), 10ng/ml Insulin-like growth factor (Peprotech, 100-11), 5µg/ml holo-Transferrin (Santa Cruz, sc-391098), 3ng/ml β-Fibroblast growth factor (R&D Systems, 4114-TC-01M), 3µg/ml heparin (Sigma-Aldrich, H3149 10KU), 0.18µg/ml hydrocortisone (Sigma-Aldrich, H0135-1MG), 10nM endothelin-1 (R&D Systems, 1160/100U), and 1x Penicillin/Streptomycin (Gibco, 15140122).

#### Targeted exon sequencing

Targeted exon sequencing was performed using a custom anchored multiplex PCR panel from ArcherDX (IDT), which was described previously^13^. DNA was extracted from cell pellets using the QIAMP DNA mini kit (Qiagen, 56304), and 150ng of DNA used for library generation according to the Archer VariantPlex HS/HGC protocol for Illumina sequencing. Library DNA was quantified with the KAPA library quantitation (Roche, KR0405) prior to sequencing on an Illumina NovaSeq X. All FASTQ files were uploaded to and analyzed on the Archer Analysis Unlimited (v7.1) website to acquire probe level CNV data. CNV were normalized to a control dataset containing eight foreskin samples which were sequenced and analyzed on the same workflow.

#### Comparison of CNV in paired patient tumors from published whole exome sequencing data

Bland-Altman plots were used to analyze CNV outliers on published targeted exon sequencing data^13^ and copy number segment data from whole exon sequencing^12,14^. To harmonize WES copy-number segments, all segment start and end positions from the paired samples were combined to define a common set of genomic breakpoints, which were used to subdivide the genome into the smallest shared intervals. Each interval was assigned the copy-number value of the original segment spanning that interval for each tumor. Intervals not covered by a reported segment were assigned a copy-number value of 1. CNV were compared between tumors that both arose from the same patient and were analyzed using the same sequencing method. Graphpad Prism version 11.1.0 was used to determine the 95^th^ percentile confidence intervals. Segments were classified as chromosome gain/loss when all outlying segments were located outside of only one confidence interval, and a complex SV/CNV event was called when nearby segments were split outside both confidence intervals. For targeted exon sequencing, more than half of probes for a gene must have fallen outside the 95^th^ percentile confidence interval for all probes within the gene to be tested for significance by Mann Whitney U against all remaining probes in the dataset. WES segments were tested for significance if two or more segments were outside the 95^th^ percentile confidence interval. All segments within and adjacent to the SV/CNV event were compared to all other WES segments using Mann Whitney U. Since WES segment data from Basurto-Lozada, *Nature*, 2026^14^ was not annotated for CAS, a filter for highly amplified CNV was set to CN≥5 and regions with complex CNV changes required at least 3 CN state changes where one was at least CN≥4.

#### OncoScan CNV array

Extracted genomic DNA (1µg) from cell culture (Qiagen DNeasy Blood & Tissue Kit, 69504) or PDX FFPE blocks were analyzed at ARUP Laboratories using their clinically-available FFPEARRAY assay (ARUP Test #3004275). CEL files were received and used to generate hg19 aligned OSCHP files that were baselined according to karyotype, if available, or baselined to a likely diploid chromosome arm using Thermo Fisher’s Chromosome Analysis Suite v4.5.0.34. SV/CNV loci were called according to criteria listed below:

Required CNV array SV/CNV locus criteria:

1. Localized high amplification spikes on whole genome view Log2R tracing (**Extended Data Figure 8b**)
2. Chromosome view of individual high amplification peaks and plateaus comprise <1/3 of the chromosome arm (**Extended Data Figure 8g-h**)
3. Region cannot be better classified as another common event (**Extended Data Figure 8d**)

Supporting criteria (requires at least 1):

1. Area of high amplification flanked by CN loss or LOH on at least one side (**Extended Data Figure 8c**, **8g-h**)
2. Non-Gaussian peaks on Log2R tracing

#### Karyotyping and metaphase spread chromosome enumeration

Cells were arrested in metaphase by incubating cells for 30 minutes in media spiked with 0.5µg/ml of colchicine (colcemid, Gibco, 15210-040). Media was collected, and cells were lifted with TrypLE (Gibco, 12605-010). After centrifuging at 300g for 7 minutes, the cell pellet was treated with hypotonic 0.075M KCl (Thermo Fisher Scientific,10575090) for 7 minutes before centrifuging at 300g for 7 minutes. The cell pellet was then fixed with a 3:1 methanol to glacial acetic acid solution for 15 min before storage at 4°C in fresh fixing solution.

Metaphase spreads were created by dropping the fixed cell suspension onto glass slides rinsed with deionized water. Once dried, the slides were either examined immediately by phase contrast or stained with a 1:1000 Hoechst 33342 solution (Thermo Fisher Scientific, 62249) for 1 minute prior to rinsing in deionized water. Both phase and Hoechst immunofluorescence images were collected on a Nikon Ti-E inverted microscope (Nikon, USA). Chromosome quantification was performed using the multipoint tool in ImageJ v1.54r with up to 200 chromosomes counted.

For Giemsa-staining, banding, and karyotyping, the remainder of the fixed metaphase-arrested cell pellets were shipped to Karyologic Inc (Durham, NC, USA) or to ARUP Laboratories (Salt Lake City, UT, USA). Karyologic Inc provided a detailed list of cytogenetic aberrations present within each analyzed metaphase spread that was used for phylogenetic analysis in R (Ape package, Manhattan method).

Cytogenetic data from other tumor types were acquired from the Mitelman Database (Mitelman Database of Chromosome Aberrations and Gene Fusions in Cancer (2026). Mitelman F, Johansson B, Mertens F and Paulsson K (Eds.), https://mitelmandatabase.isb-cgc.org)^19^.

#### Nuclear ploidy by flow cytometry

PDX tumor FFPE blocks were sent to ARUP Laboratories for testing on their clinically available “DNA cell cycle analysis – Ploidy and S-phase” (ARUP test code 0095155, DNA MISC). We received .fcs files with the presumptive G0/G1 peak (P1) set to 50,000 on the propidium iodide channel. All subsequent analysis and quantification were performed using FlowJo v10.10.0 (Waters Biosciences, Ashland, OR, USA).

#### Immunofluorescence and nuclear abnormality quantification

Primary human melanocytes, CM, and AM cells were seeded onto clear poly-L-lysine-coated glass coverslips in 6-well plates at a density of 300,000 cells per well. After 48 hours of incubation, cells were washed with HBSS and fixed with 4% paraformaldehyde at room temperature for 20 minutes. Coverslips were then washed with HBSS, permeabilized with 0.25% triton-x in PBS for 10 minutes at room temperature, and washed with PBS. Cells were stained with ActinRed™ 555 ReadyProbes™ Reagent (Invitrogen, R37112) according to the manufacturer’s instructions for 30 minutes at room temperature in the dark, washed with PBS, and mounted on slides using DAPI Fluoromount-G® (Southern Biotech, 0100-20). Images were acquired on a Nikon Ti-E inverted microscope (Nikon, USA) using identical exposure times for all samples and the same magnification settings.

Nuclear abnormalities were quantified by counting the total number of DAPI-positive DNA bridges across the entire coverslip. The number of DNA-containing anaphase bridges per cell was then calculated using the average number of cells per field of view and the total number of fields of view per coverslip. Anaphase bridge numbers were normalized to NHM and plotted as fold change. For nuclear buds and micronuclei, 30 random fields of view per coverslip were imaged. Nuclear buds were defined as DAPI-positive circular bodies attached to the nucleus. Micronuclei were defined as DAPI-positive circular bodies not attached to the nucleus but present within the same cell based on actin staining. The number of nuclei, micronuclei, and nuclear buds was quantified per field of view. The frequency of nuclear abnormalities (combined micronuclei and nuclear buds) was calculated as the ratio of nuclear abnormalities to the total number of nuclei.

#### Bulk tumor whole genome sequencing and JaBba analysis

Dissociated and mouse-depleted PDX tumor cells underwent DNA extraction with the QIAMP DNA mini kit (Qiagen 56304) and library preparation with NEBNext Ultra II DNA PCR-free Library Prep Kit (New England Biolabs E7410S). Libraries were prepared and sequenced on a NovaSeq X at the HCI High Throughput Genomics core facility. JaBba was built from source along with its dependancies including Gurobi, fragCounter, and SvABA (https://github.com/mskilab-org/JaBbA, https://www.gurobi.com, https://github.com/mskilab-org/fragCounter, https://github.com/walaj/svaba). Each application was run using default parameters and the GRCh38 GIABv3 human reference. For the HCI-AM087 sample 25595X2, the JaBba modeling parameters were relaxed to enable convergence (--tilim 120000 --epgap 1e-2). Where a normal was required, a public WGS from CEPH individual NA12878 was downloaded from Illumina’s BaseSpace repository (NovaSeqX-TSPF-NA12878-UDI-H09) and processed as the AM WGS samples.

The JaBba output was visualized with the gGnome R package (https://github.com/mskilab-org/gGnome).

#### Single cell whole genome sequencing

HCI-AM087 and HCI-CM019 PDX tumors were dissociated and depleted of mouse stroma (see above) before being cryopreserved in 90% FBS and 10% DMSO. A vial from each tumor was sent to BioSkryb Genomics (Durham, NC, USA) for viable cell sorting, individual cell library preparation using ResolveDNA primary template-direct amplification^43^, and low-depth quality control sequencing. Additionally, five CM (24316x1-5) and fifteen AM (24316x6-20) single cell libraries were chosen for 30x WGS on an Illumina NovaSeq X at the HCI High Throughput Genomics resource. These sequences were aligned to GRCh38/hg38 for downstream analysis.

#### Structural variant calling, filtering, quantification, and mapping

SVs were called on all reads using Manta (v1.6.0) and the GRCh38 reference genome (GIABv3, no alt, masked GRC decoys) to generate candidate structural variant calls for each sample. Candidate SVs from all samples were aggregated into a unified dataset. During aggregation, the SVTYPE was extracted from the INFO column, and absolute genomic positions were calculated to facilitate cross-chromosome analysis. To focus on high confidence breakends originating within specific target genomic regions, the ALT field was parsed to determine the target chromosome and position. The number of supporting read pairs for each breakend was extracted from the BND_PAIR_COUNT field in the INFO column. The resulting dataset was then strictly filtered to retain only structural variants supported by a minimum of 5 read pairs.

To reconstruct the phylogenetic relationships between samples based on the stringent SV filtering, a presence/absence matrix of the filtered structural variants across all samples was constructed. A pairwise distance matrix was then computed using the non-Euclidean Jaccard distance metric to quantify the dissimilarity of shared structural variant profiles between samples. Hierarchical clustering was performed on the resulting distance matrix using the unweighted pair group method with arithmetic mean (UPGMA, or ‘average’ linkage). Finally, the clustered relationships were visualized as a dendrogram (phylogenetic tree) and annotated with the specific shared breakends at each branching node to trace the evolutionary acquisition of the structural variants.

To evaluate the frequency of structural variant sharing within sample subpopulations, a combinatorial sampling analysis was performed across the whole genome. Breakend variants from all samples were first canonicalized to ensure consistent coordinate representation and subsequently clustered into 100-base-pair genomic bins to account for minor breakpoint calling variations. Samples were strictly partitioned into two cohorts: Group A (samples X1–X5) and Group B (samples X6–X20). To assess variant sharing at the triplet level, every possible unique combination of three samples (triplets) was generated within each respective group. For each triplet, the total number of whole-genome breakend clusters completely shared among all three constituent samples was calculated. The resulting distributions of shared event counts for Group A triplets versus Group B triplets were visualized as superimposed swarm and box plots. Statistical significance of the difference in shared breakpoint frequencies between the two groups was determined using a two-sided Mann-Whitney U.

Breakpoint data aggregation, filtering, and statistical analyses were performed using Python 3, utilizing the pandas and numpy libraries for data manipulation. Hierarchical clustering and distance matrix computations were conducted using the scipy.cluster.hierarchy and scipy.spatial.distance modules. Non-parametric statistical comparisons between groups, specifically the two-sided Mann-Whitney U, were performed using the scipy.stats module. Boxplots, swarm plots, and dendrograms were generated using the matplotlib and seaborn libraries. Finally, circular genome visualizations depicting the structural variant breakpoints were generated using the pycirclize Python package.

#### FBI identification and mapping at GAB2 locus

For FBI identification and mapping, LL, RR, and RL pairwise reads were extracted using the SAMtools command ‘samtools view -b -F 1294 -T’ and placed into a BAM file. Using the LL, RR, and RL enriched BAM files, intrachromosomal FBI were identified at the GAB2 locus by identifying ≥2 non-identical reads in ≥2 cells that contain the same breakpoint and soft read profiles. BLAT was used to identify the coordinates and distance of the other junction using a minimal score cutoff of 750. A composite of all FBI and FBI within a sample of cells were visualized in Integrative Genomics Viewer, and a phylogenetic relationship tree was generated in R (Ape package, Manhattan method) to visualize the differences in FBI fingerprint between individual cells.

#### RNA-seq and gene fusion identification

The HCI High Throughput Genomics Core performed all quality control, library preparation, and sequencing for RNA isolated from flash-frozen PDX tumors. Quality control and quantifications were performed using an Agilent High Sensitivity RNA ScreenTape Assay (Agilent, 5067-5579) on an Agilent 2200 TapeStation. The NEBNext Ultra II Directional RNA Library Prep (New England Biolabs, E7760L) and rRNA Depletion Kit v2 (E7400X) were used to generate libraries according to the manufacturer’s protocol. Sequencing of the libraries was performed on an Illumina NovaSeq X to acquire approximately 35 million reads per sample.

Alignment and downstream bioinformatic analysis was performed by the HCI Cancer Bioinformatics Shared Resource (CBI). Mouse sequences were removed using a GRCh38-GRCm39 combined FASTA reference. This reference was indexed with STAR and used to align each RNAseq dataset. Human sequences were extracted by creating a SAMTools filter for mapping quality threshold of 13 and for reads that uniquely mapped to only human sequences. This process was supported using a containerized Snakemake workflow (https://github.com/HuntsmanCancerInstitute/Workflows/tree/master/Hg38RunnerWorkflows2/RnaAlignQC/HumanPdx.)

A list of putative gene fusions was identified using STAR-Fusion by the CBI. To be considered as positive fusion, there must be ≥10 junction read count, ≥2 spanning fragment counts, ≥0.5 fusion fragments per million total reads, and >0.1 left and right break entropy.

#### Statistics

If not otherwise mentioned above, all statistical tests were performed in Graphpad Prism v11.1.0.

#### Data availability

OncoScan CNV .CEL and .OSCHP files are located within Source Data 3. PDX RNA-sequencing data and single-cell and bulk tumor whole-genome sequencing FASTQ files for HCI-AM087 and HCI-CM019 will be deposited in [dbGaP/GEO] under controlled access prior to publication in a peer-reviewed journal. Published WES^12,14^ and targeted exon sequencing^13^ are available through their respective journals.

#### Figure generation

All figures were generated using a combination of R 4.6.1, Matplotlib 3.10.5, python 3.11.13, Graphpad Prism v11.1.0 Adobe Illustrator 30.8.2, and Microsoft PowerPoint 365.

## Extended Data Figures and Legends

**Extended Data Figure 1:**
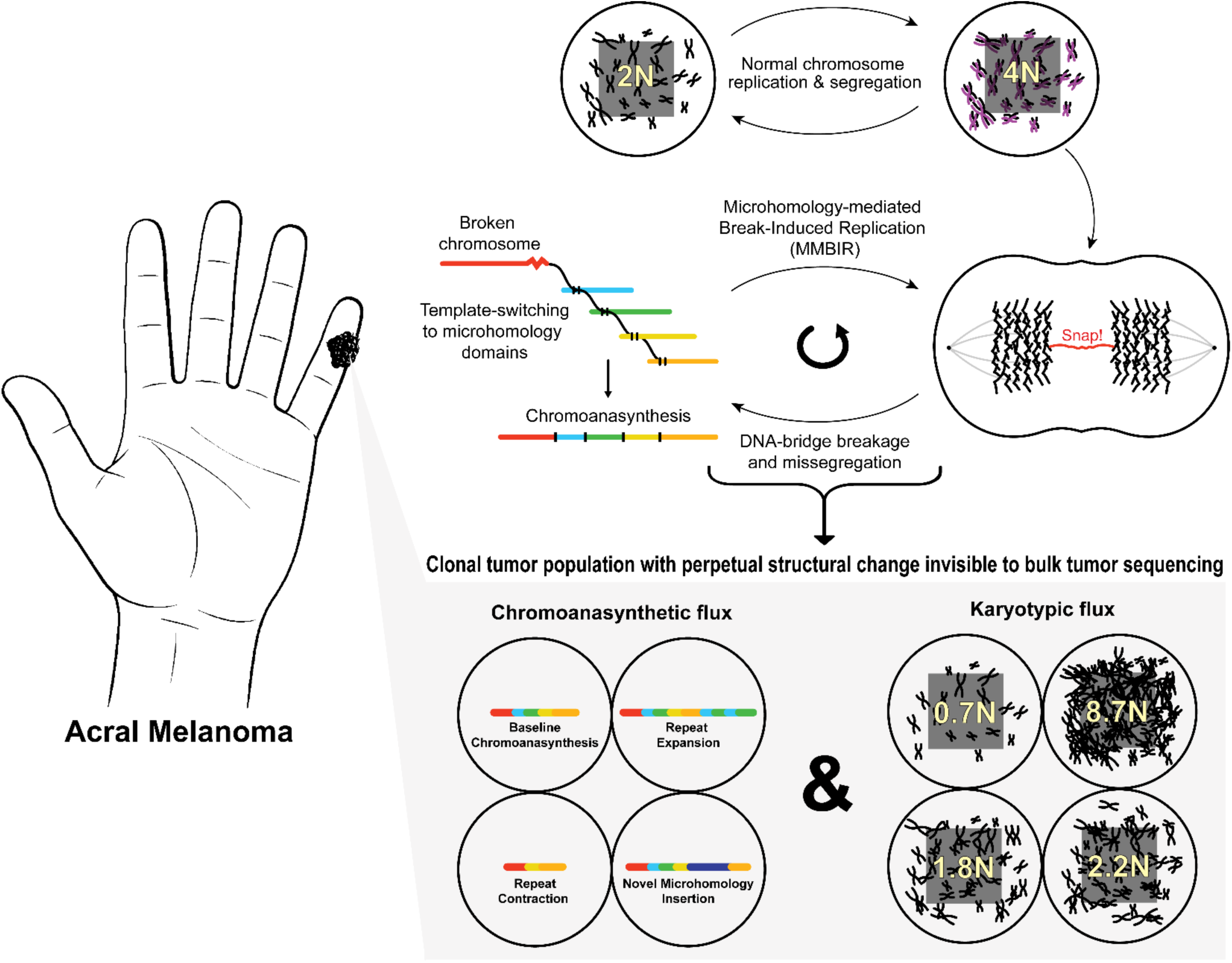
Visual Abstract. Acral Melanoma undergoes large fluctuations in chromosome numbers during cell division (Karyotypic-flux), which is associated with DNA bridge formation. In parallel, chromoanasynthesis from microhomology-mediated break-induced replication (MMBIR) DNA repair drives novel structural variations observed in all sequenced cell genomes (Chromoanasynthetic-flux).

**Extended Data Figure 2:**
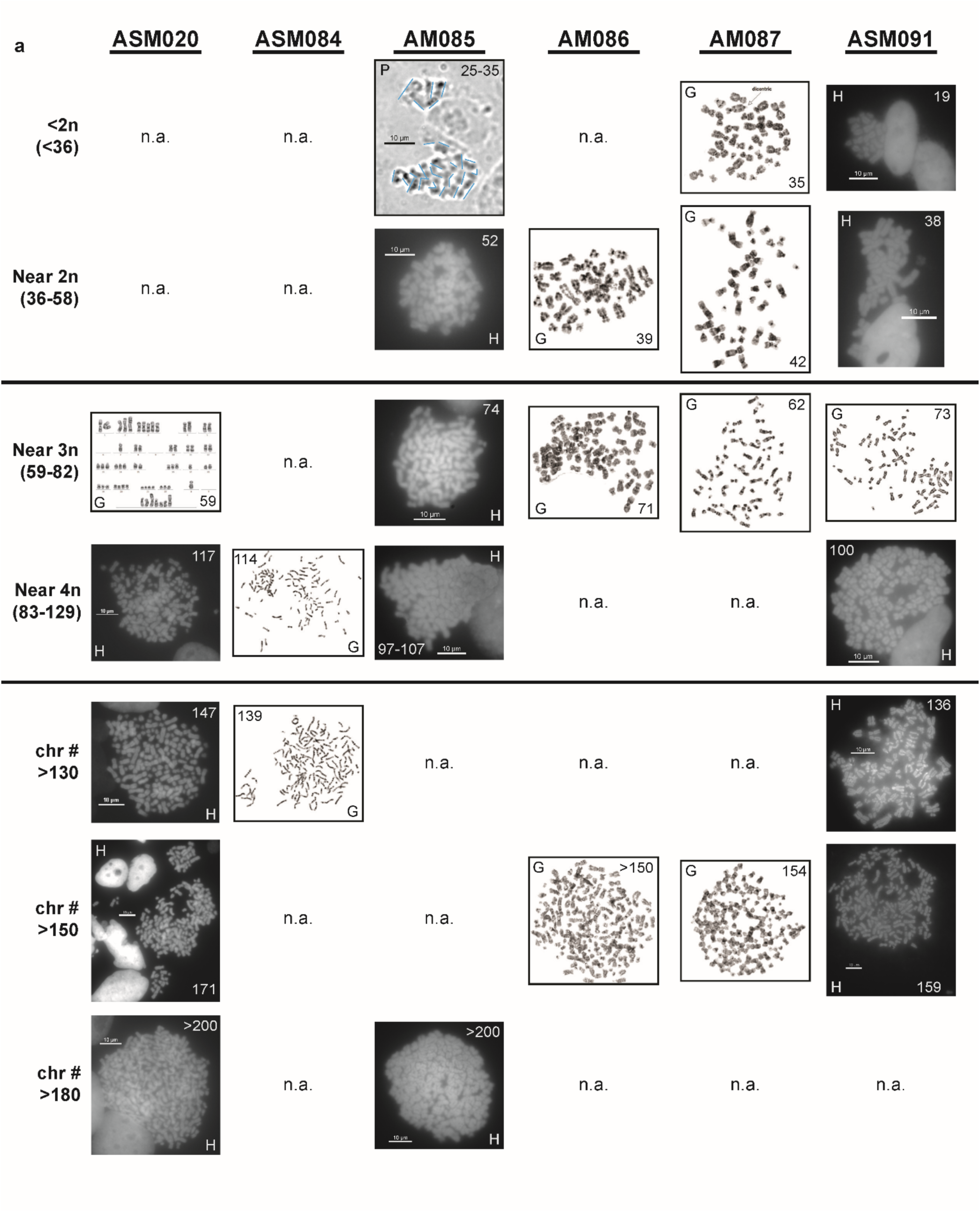
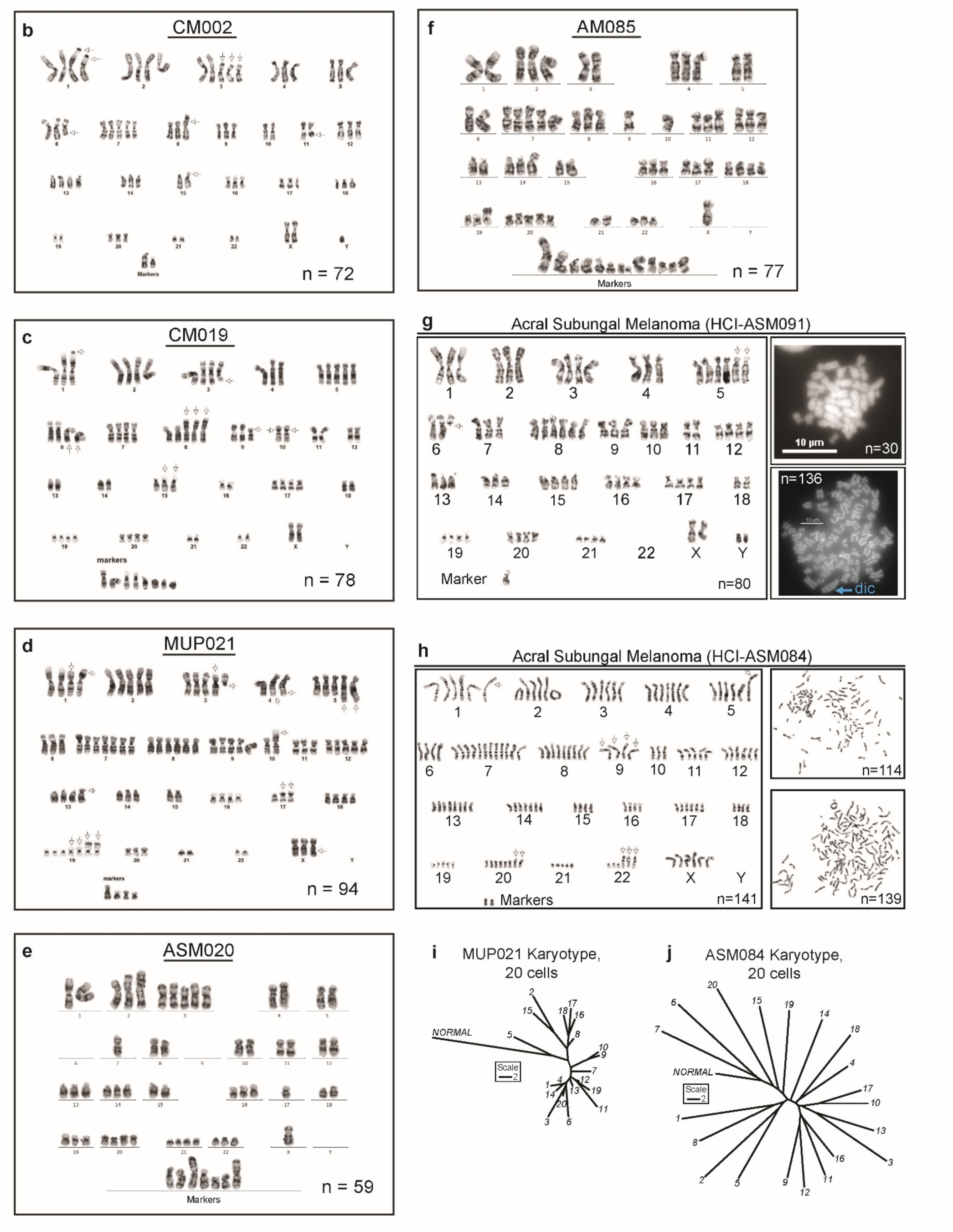
Representative metaphase spreads images and karyograms demonstrating K-flux positivity in AM tumor models. **a**, Images of metaphase spreads from six different cell cultures are arranged according to chromosome count. Three types of images are present based on technique. ‘P’ represents phase contrast, and an estimated chromosome count is provided along with blue outlines of chromosomes. ‘H’ represents Hoechst-stained metaphase spreads, and ‘G’ represents Giemsa-stained metaphase spreads or karyograms. All scale bars are 10µm, and chromosome counts are indicated on the image. Only ASM084 is K-flux negative. All other AM spreads are K-flux positive (>46 chromosome difference between maximum and minimum observed chromosome counts). **b-c,** Representative karyograms for CM and **d**, melanoma of unknown primary (MUP). **e-f,** HCI-ASM020 and HCI-AM085 had predominately low-resolution metaphase spreads. Example spreads that could be Giemsa-banded are shown. Representative karyograms and metaphase spreads for **g**, HCI-ASM091 and **h**, HCI-ASM091 also highlight K-flux. Not shown: all metaphase spreads for AM086 were too low quality for karyogram. **i-j**, Phylogenetic trees based on karyogram data show tight clustering in HCI-MUP021 and high variability in HCI-ASM084.

**Extended Data Figure 3:**
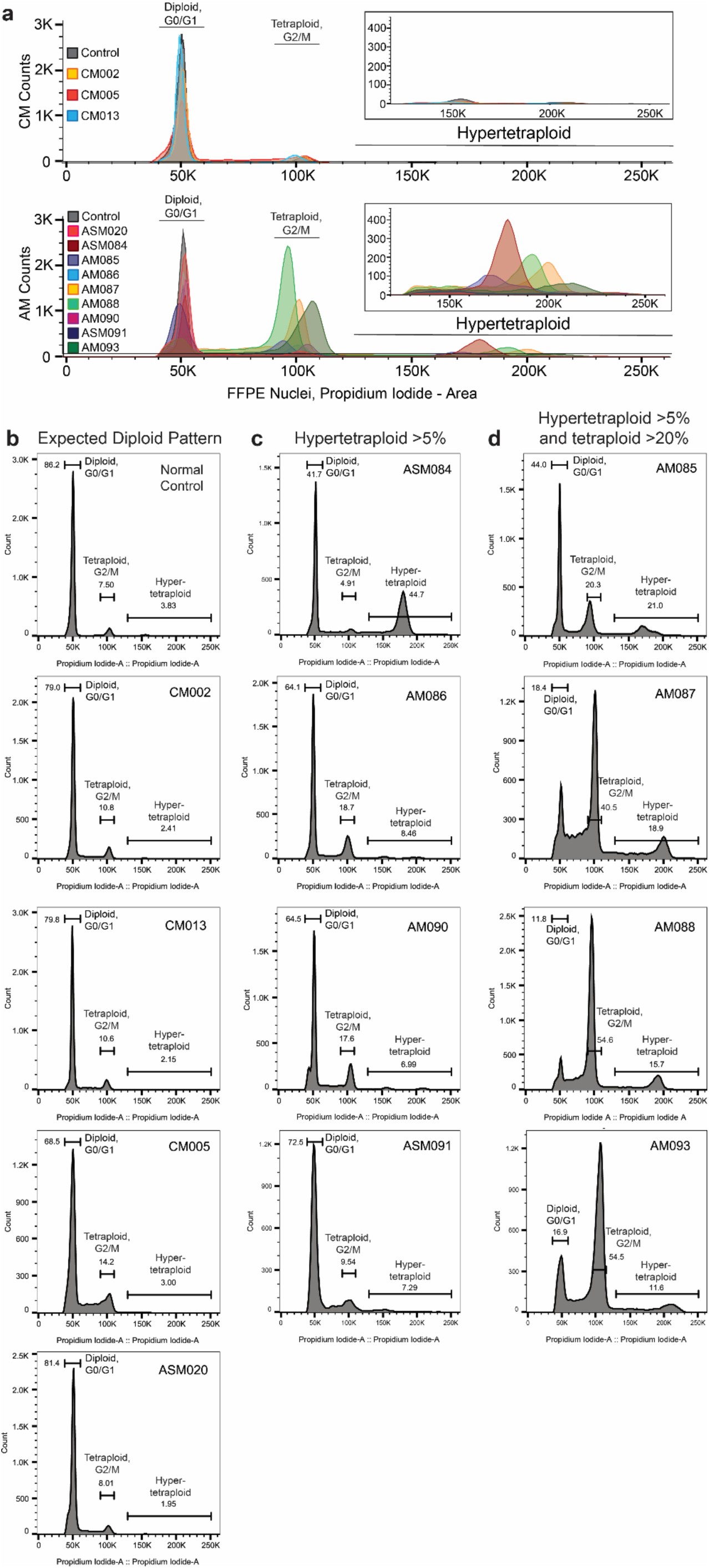
Individual nuclear flow cytometry histograms for each tumor model. **a**, The top aggregate histograms demonstrate the expected diploid G1-predominate cell cycle patterns for normal control and CM tumors. The bottom histogram showcases the variability of G2/M and hypertetraploidy dominant ploidy patterns in AM. **b**, Individual histograms for control, CM tumors, and HCI-ASM020 show expected diploid cell cycle peak distribution. **c**, Four AM tumor models exhibited an increased hypertetraploidy population of nuclei. **d**, An additional four AM tumor models had the increased hypertetraploidy population with an enriched tetraploidy or G2/M population.

**Extended Data Figure 4:**
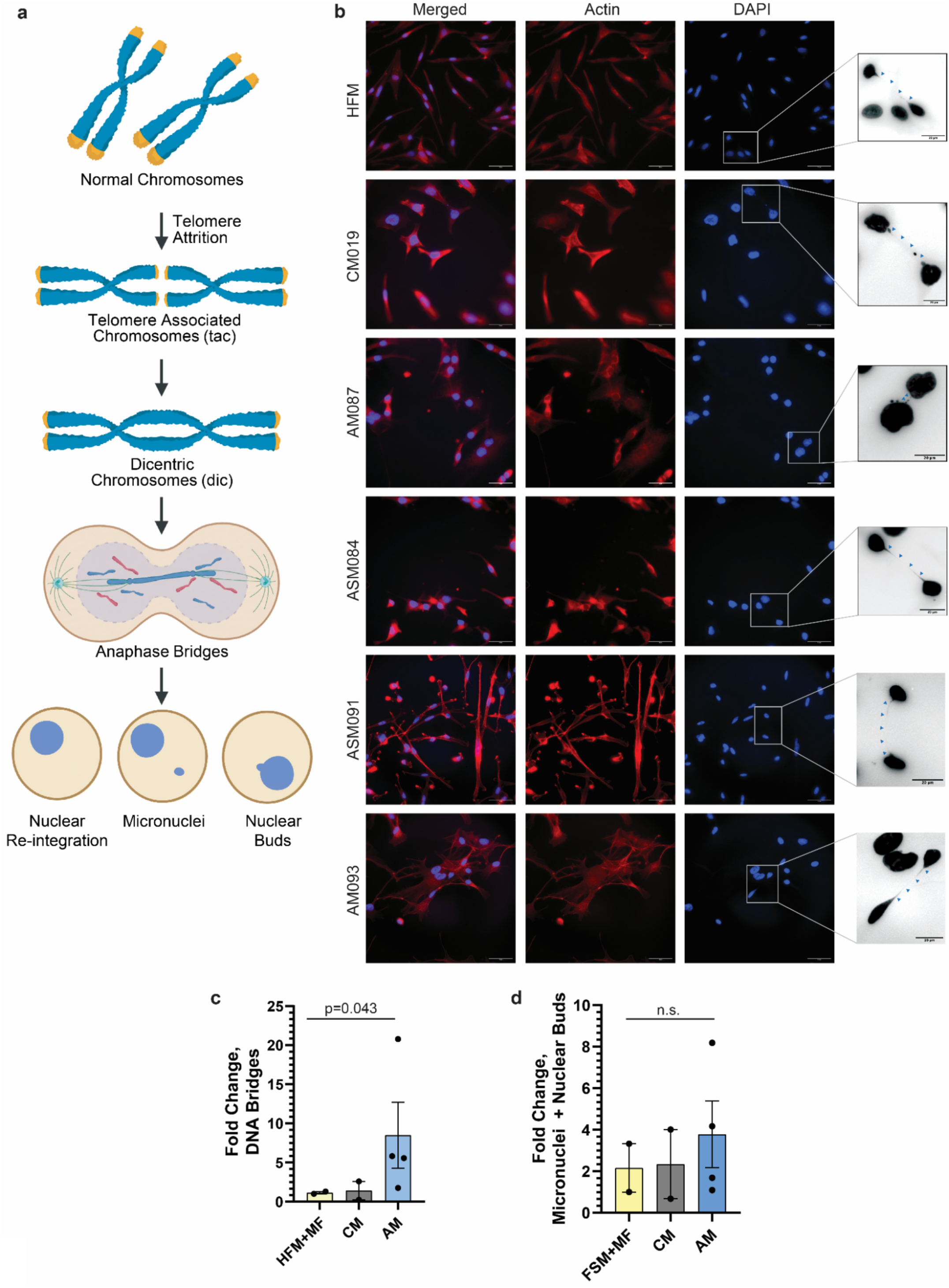
Most AM models carry dicentric chromosomes and anaphase bridges without accompanying nuclear abnormalities. **a**, Telomere attrition from cell division can lead to non-covalent telomere association chromosomes (tac) during cell division. After subsequent cell divisions, the telomere length becomes low enough to provoke end-joining repair, resulting in a covalently bound dicentric chromosome (dic). Among other mechanisms, anaphase bridges can result from dic, and the surviving cells can be left with micronuclei or nuclear buds if the late segregating mitotic DNA is not incorporated into the nucleus. Modified from BioRender. Smith, E. (2026) https://BioRender.com/vuil1kh. **b**, Anaphase bridges are resolved by DAPI and actin (Alexa Fluor 488) immunofluorescence (scale bars are 50µm). Black and white inserts are magnified color inversions of DAPI anaphase bridges with blue arrowheads highlighting bridges (scale bars are 20µm). **c**, AM models have increased DNA bridges compared to non-cancerous primary human foreskin melanocytes (HFM), mouse fibroblasts (MF), and CM cell controls (one-sided exact permutation Mann-Whitney, p=0.0429). **d**, Micronuclei and nuclear bud frequency are similar across the different cell models.

**Extended Data Figure 5:**
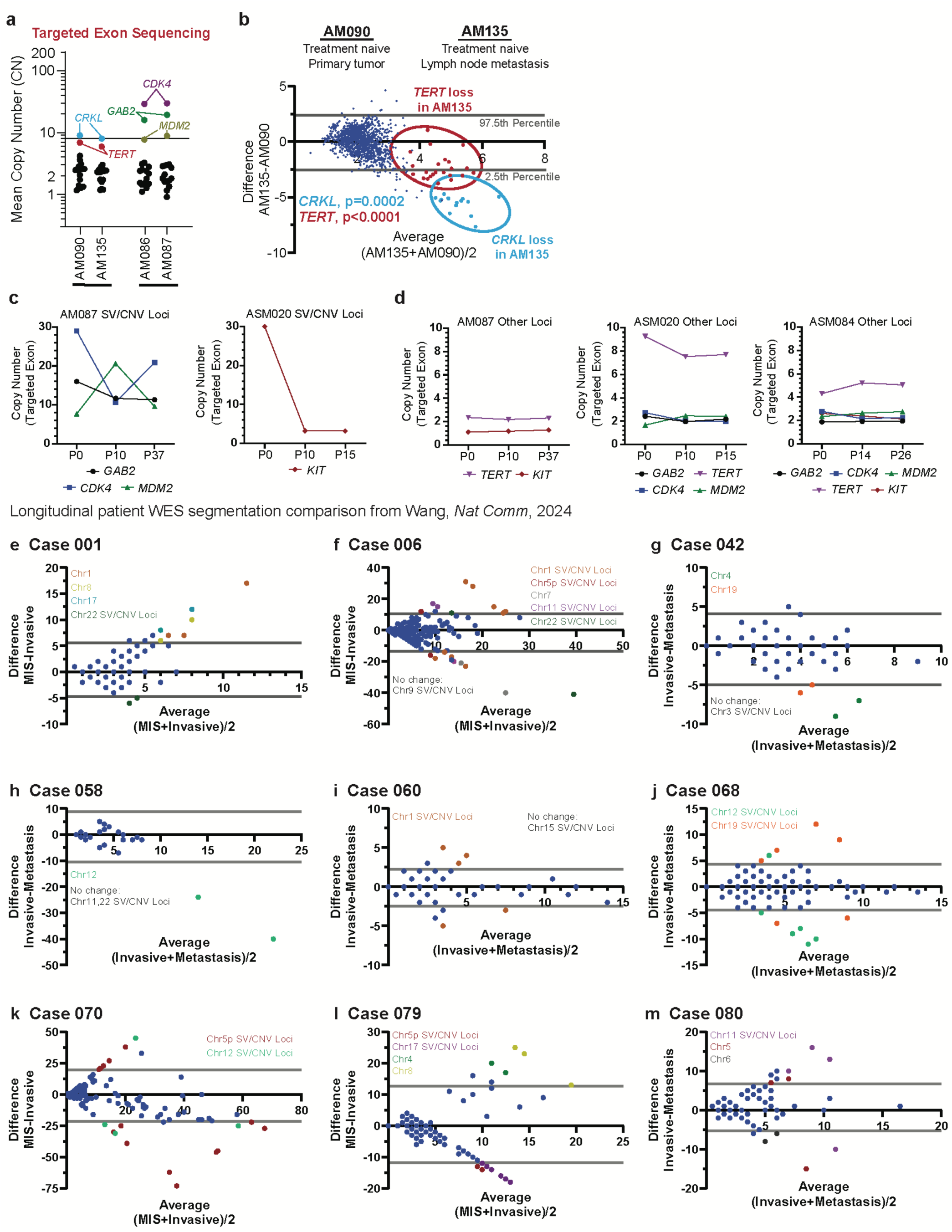
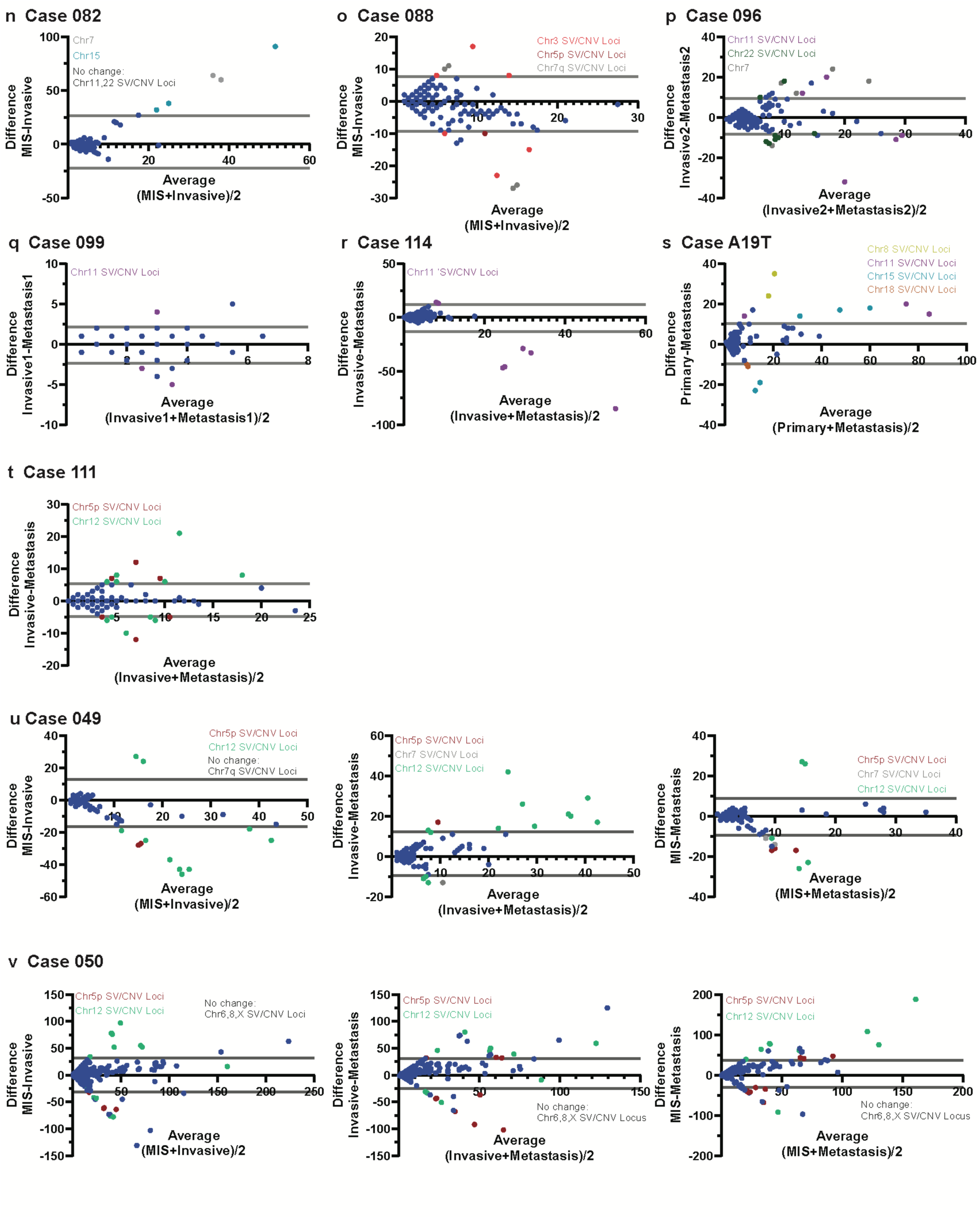
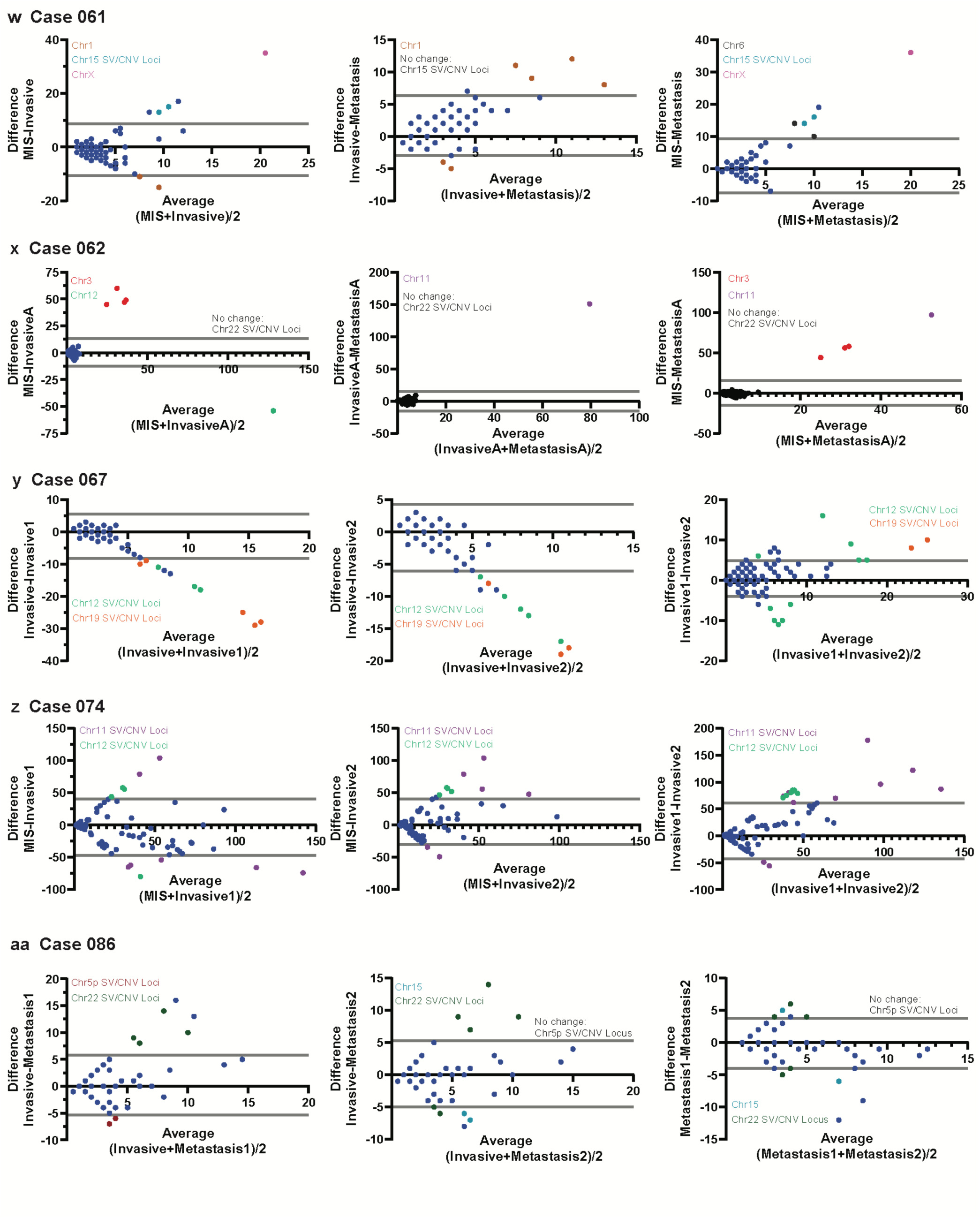
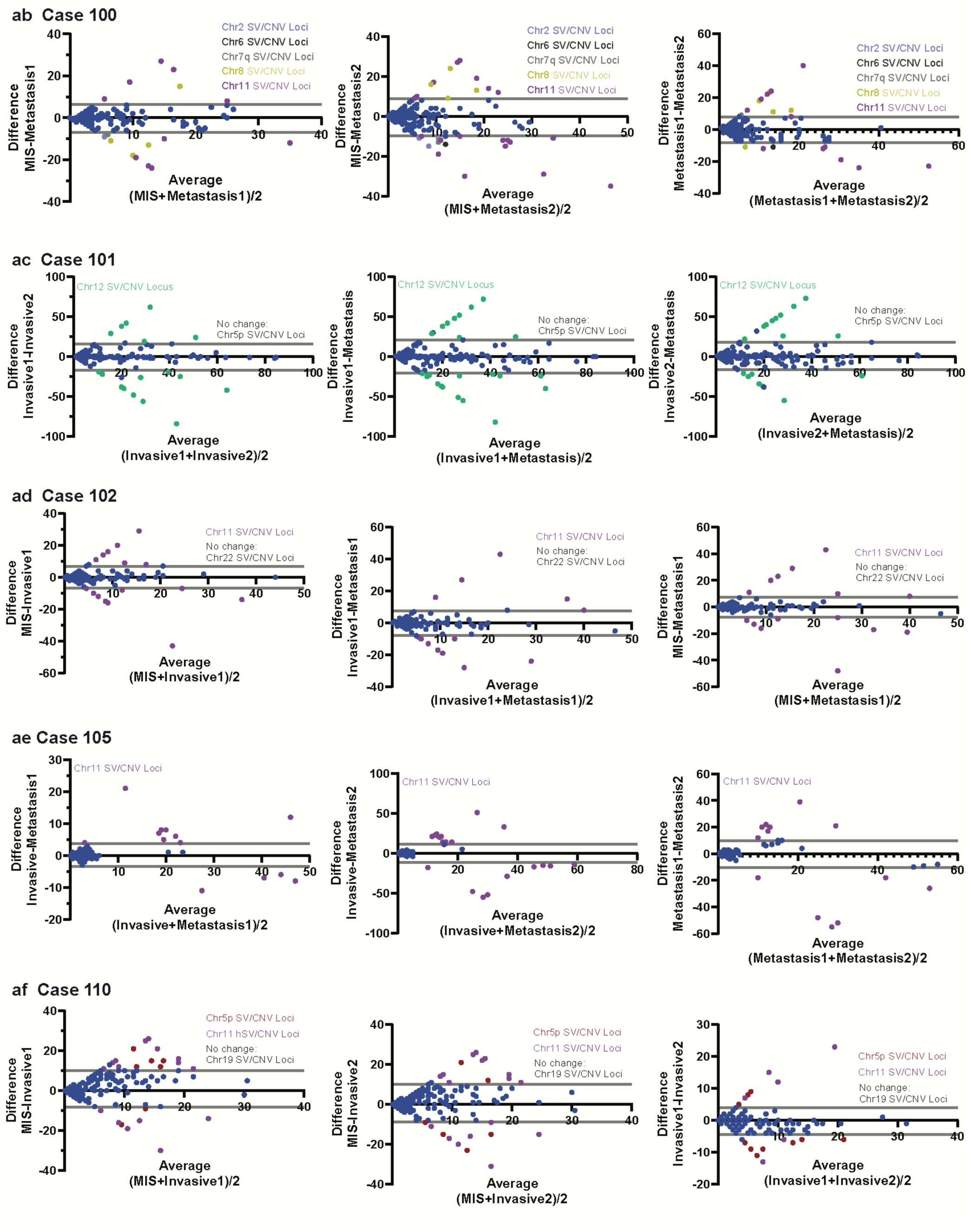
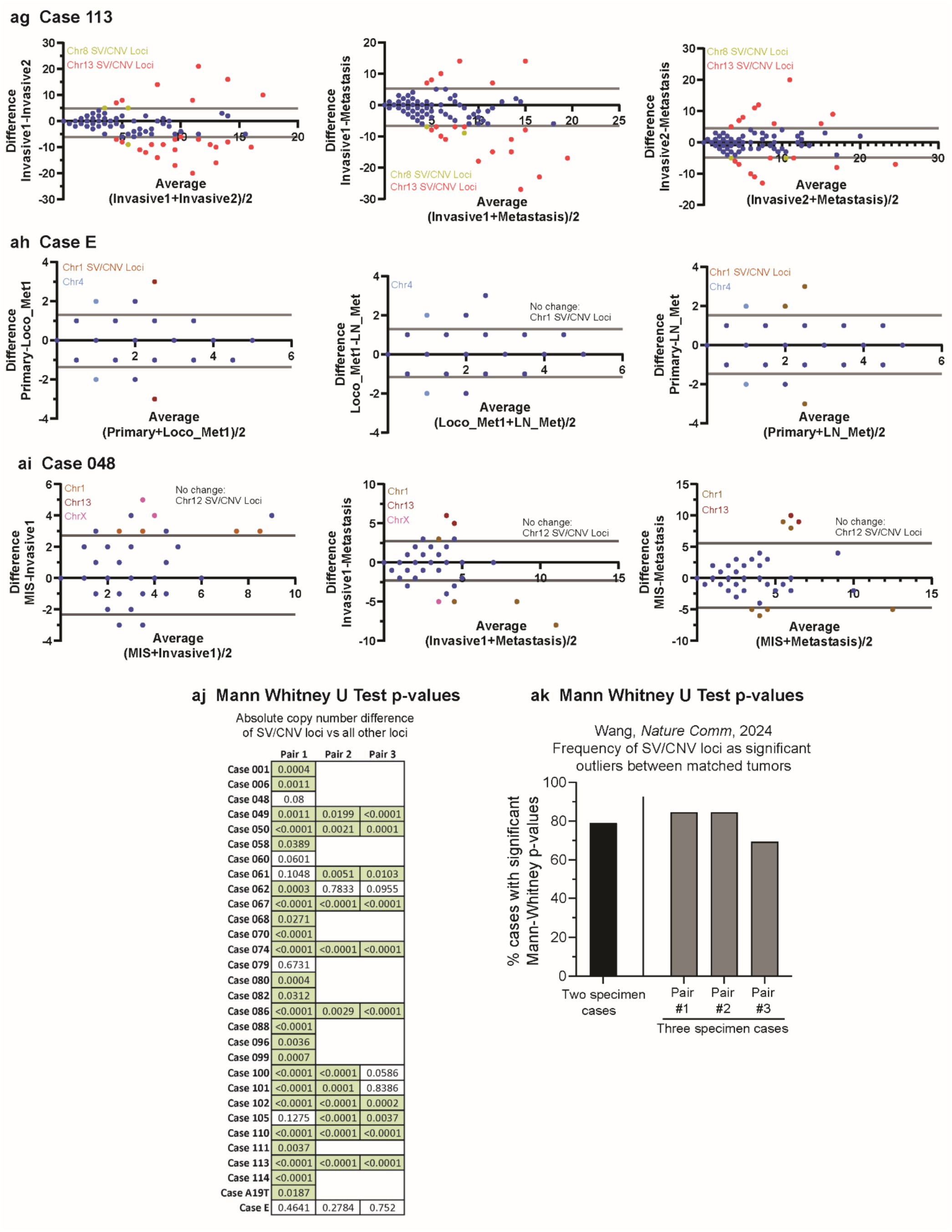
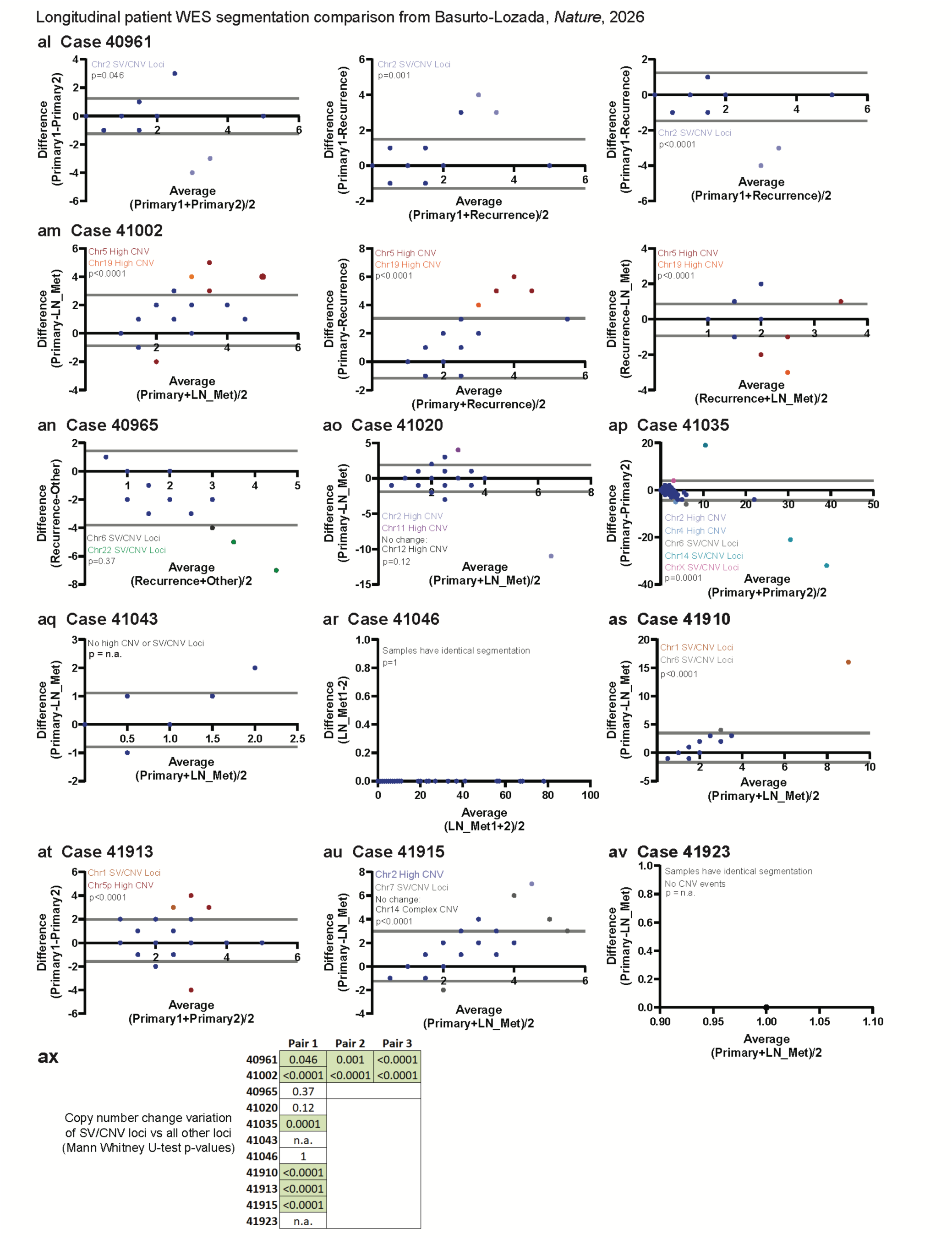
Comparison of copy number segmentation between paired tumors in PDX models, Wang, *Nat Comm*, 2024^12^, and Basurto-Lozada, *Nature*, 2026^14^. **a**, Using the targeted ArcherDX exon sequencing panel, *GAB2*, *CDK4*, and *MDM2* genes undergo an increase in CN between HCI-AM086 and HCI-AM087, and a decrease in *CRKL* and *TERT* CN between HCI-AM090 and HCI-AM135. Y-axis is log10 scale. **b**, HCI-AM090 and HCI-AM135 are patient-matched tumor progression stages collected at the same time and show a significant decrease in *TERT* and *CRKL* copy number in the HCI-AM135 (Mann-Whitney U). **c-d**, Highly amplified (CN>8) SV/CNV loci show CN fluctuation by serial targeted exon sequencing during cell culture. Loci with moderate amplification (CN<4) are more stable. Passage number is indicated on the x-axis. **e-ai**, Bland-Altman plots are used to visualize outliers between pairs of tumors from the same patient in Wang, *Nat Comm*, 2024^12^. The upper and lower 95^th^ percentile intervals are indicated by grey lines, and chromosome segments that present ≥2 times above or below the segments are colored per the key. All SV/CNV loci are included in the key, including non-outliers. **e**, Case 001 has chromosome gain/loss change. All outliers from the same chromosome are present outside of the same 95^th^ percentile confidence interval. **f**, Case 006 also has chromosome gain/loss in Chr7, but all other outliers are split between the upper and lower confidence intervals, consistent with new complex SV/CNV events. **g-t**, Remainder of cases comparing two different tumors. **u-ai**, For cases with 3 or more tumors, a selection of three tumors was compared. When possible, different tumor states were compared from the following list: MIS, primary tumor (primary), invasive, metastatic, lymph node metastasis (LN_Met), and locoregional metastasis (Loco_Met). **aj-ak**, The majority (79.3%) of paired tumor comparisons from Wang, *Nature Comm* 2024 were significant by Mann Whitney U (p-values in green boxes are significant). **al-av**, The same analysis was performed on segmented WES datasets from Basurto-Lozada, *Nature*, 2026^14^. Since this population was not enriched or annotated for SV/CNV loci, clusters of potential SV/CNV loci and highly amplified CNV segments were annotated. **al-av**, Where SV/CNV loci and High CNV events occurred, they represent outliers in CNV change between paired tumors. Some paired cases had no CNV clusters or high events (**aq**), and others had identical segmentation between paired tumors (**ar**, **av**). **ax**, Ten of 13 pair comparisons with CNV events had significant p-values as indicated by green highlighted cells (Mann Whitney U).

**Extended Data Figure 6:**
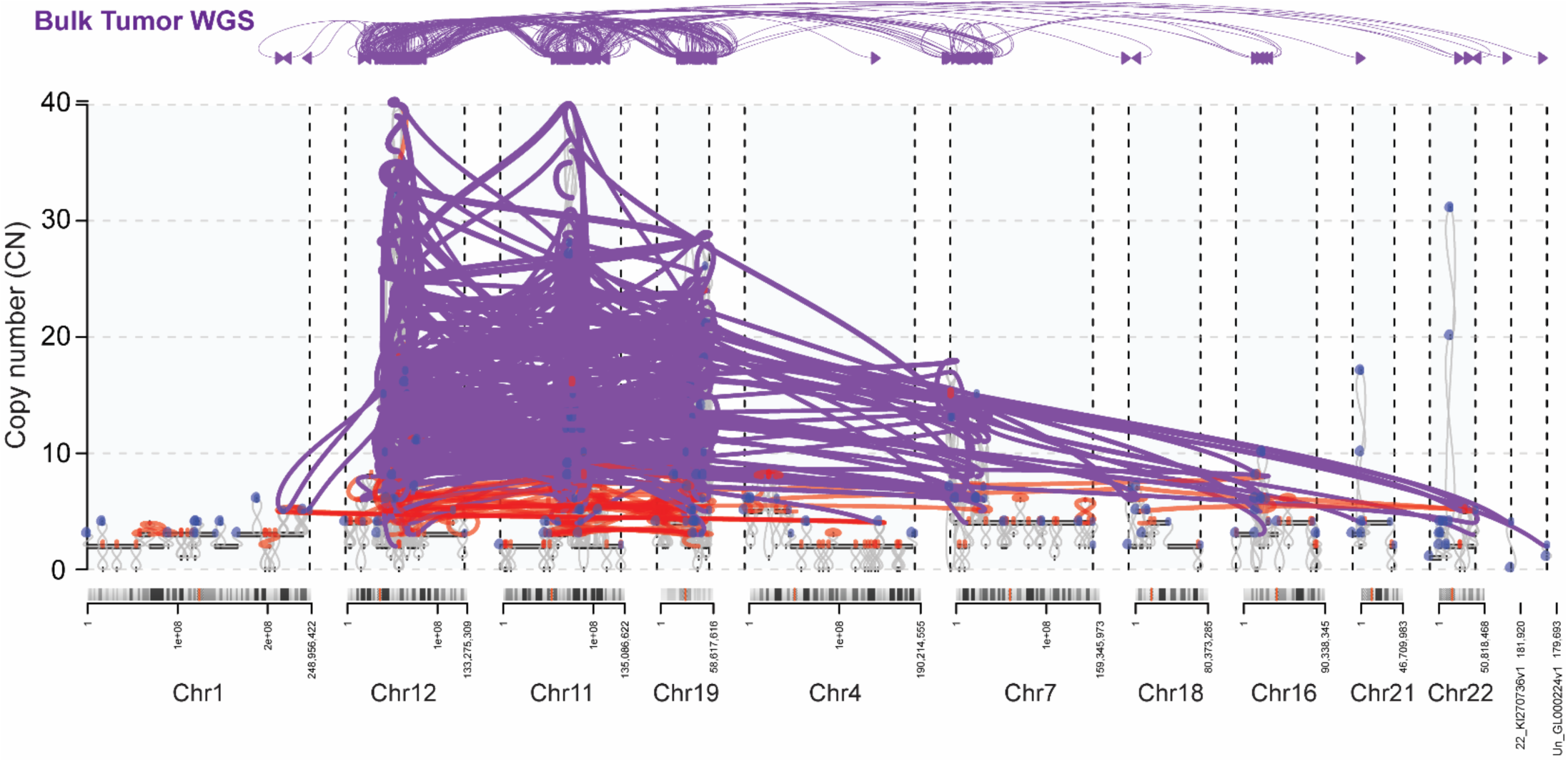
AM SV/CNV loci match the tyfonas pattern from Hadi, *Cell*, 2020^18^. JaBba and gGnome were used to map SV/CNV loci (purple) from bulk-tumor WGS in HCI-AM087. The density and orientation of complex connections is shown in the top graph (purple). The middle graph plots the type of connection between CN state changes, where grey lines represent CN changes to the adjacent reference genome, red represents alternative connections to non-adjacent sequences, and purple represents clusters of quasi-reciprocal alternative connections that are common in complex events such SV/CNV loci. Blue hooks are loose ends where an adjacent region could not be determined. The full banding of each chromosome involved in the SV/CNV event is shown above the chromosome coordinates with orange bars marking the centromeres.

**Extended Data Figure 7:**
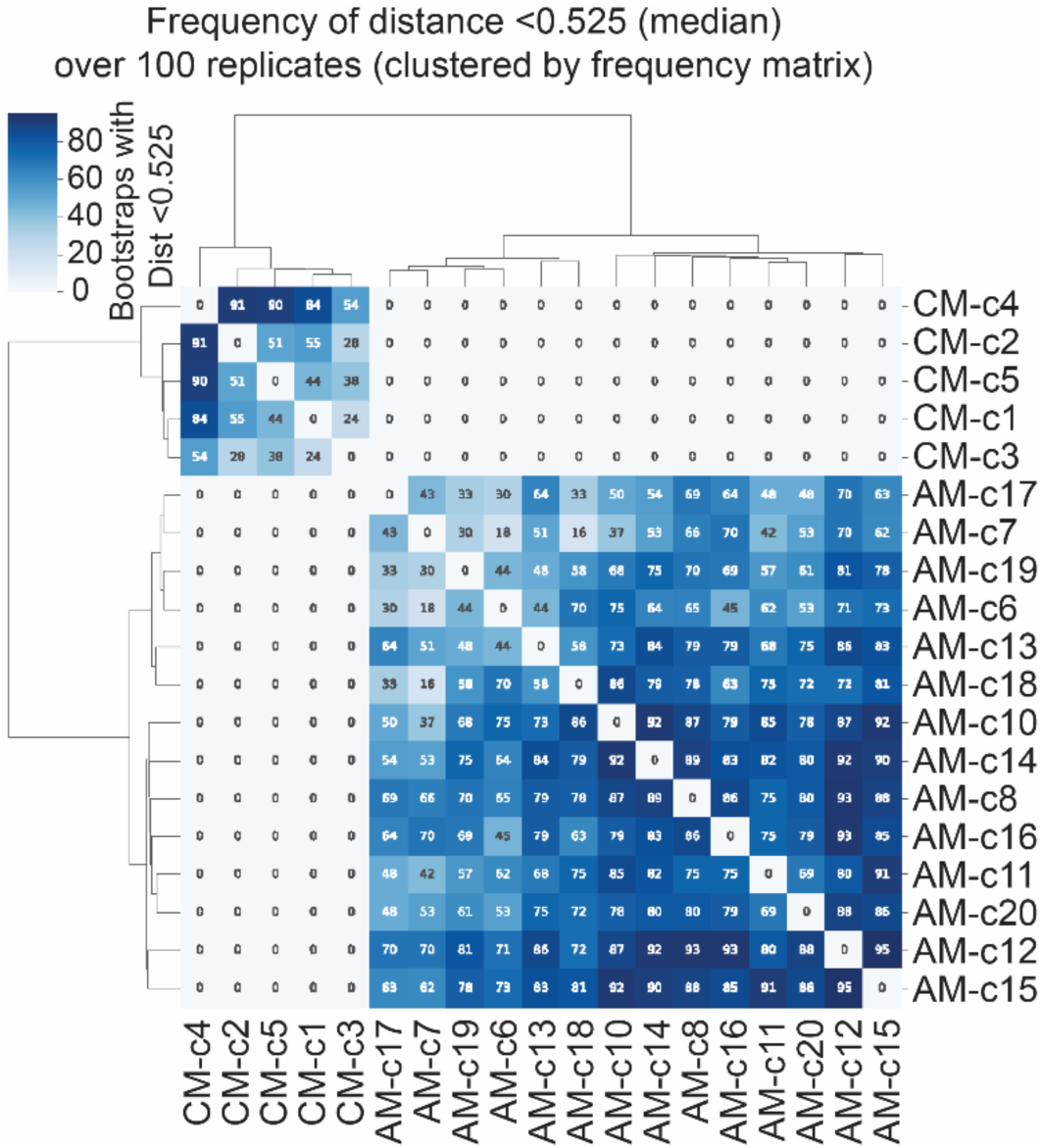
SV breakpoints independently cluster within HCI-AM087 and within HCI-CM019. Heatmap showing high similarity between five HCI-CM019 cells (CM-c1 through c-5) and 14 HCI-AM087 cells (AMc-6 through c-20). The values within each square are the Jaccard distance generated from 100 bootstrap replicates where 1000 random 10kb genomic windows containing SV were sampled.

**Extended Data Figure 8:**
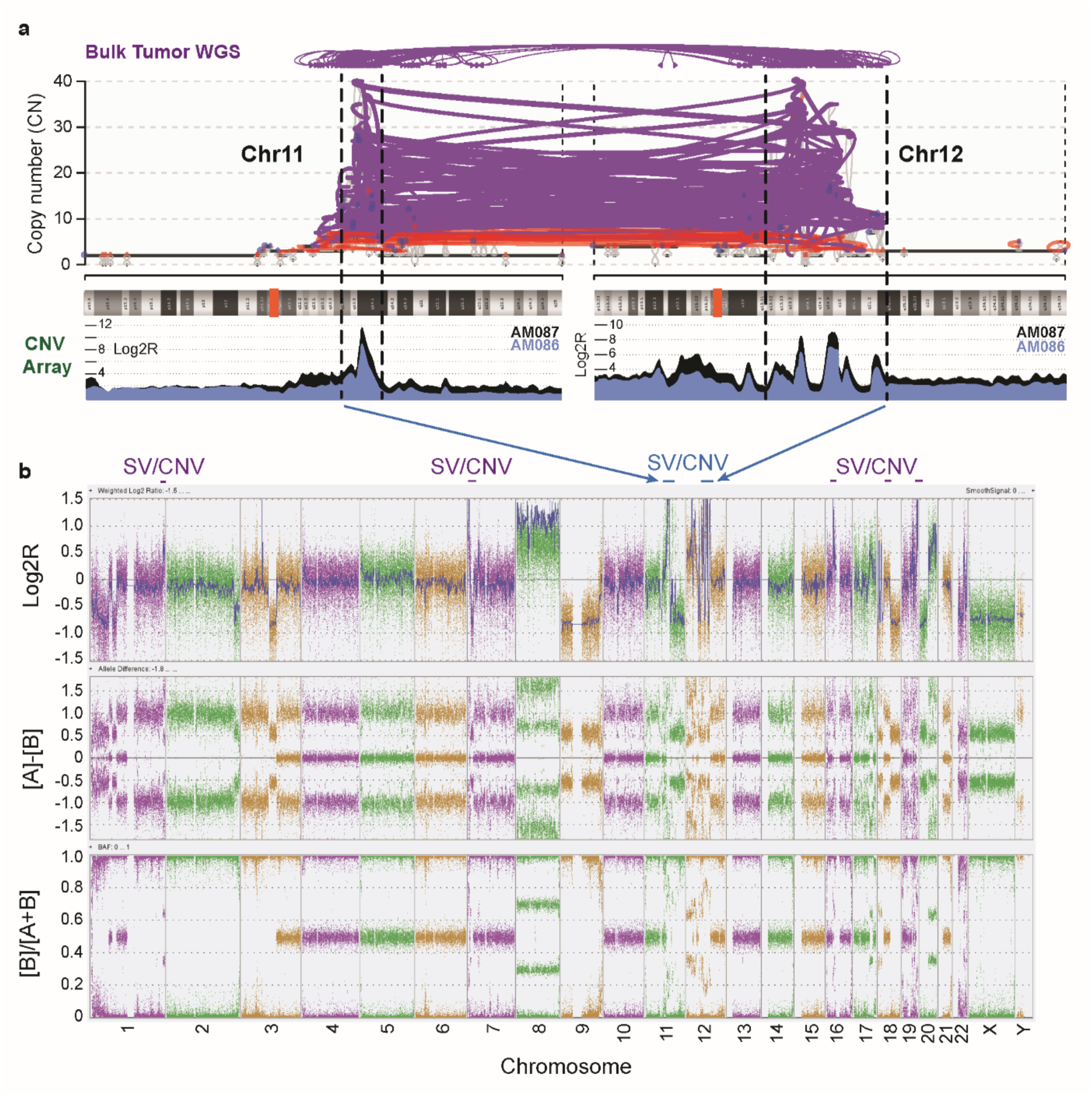
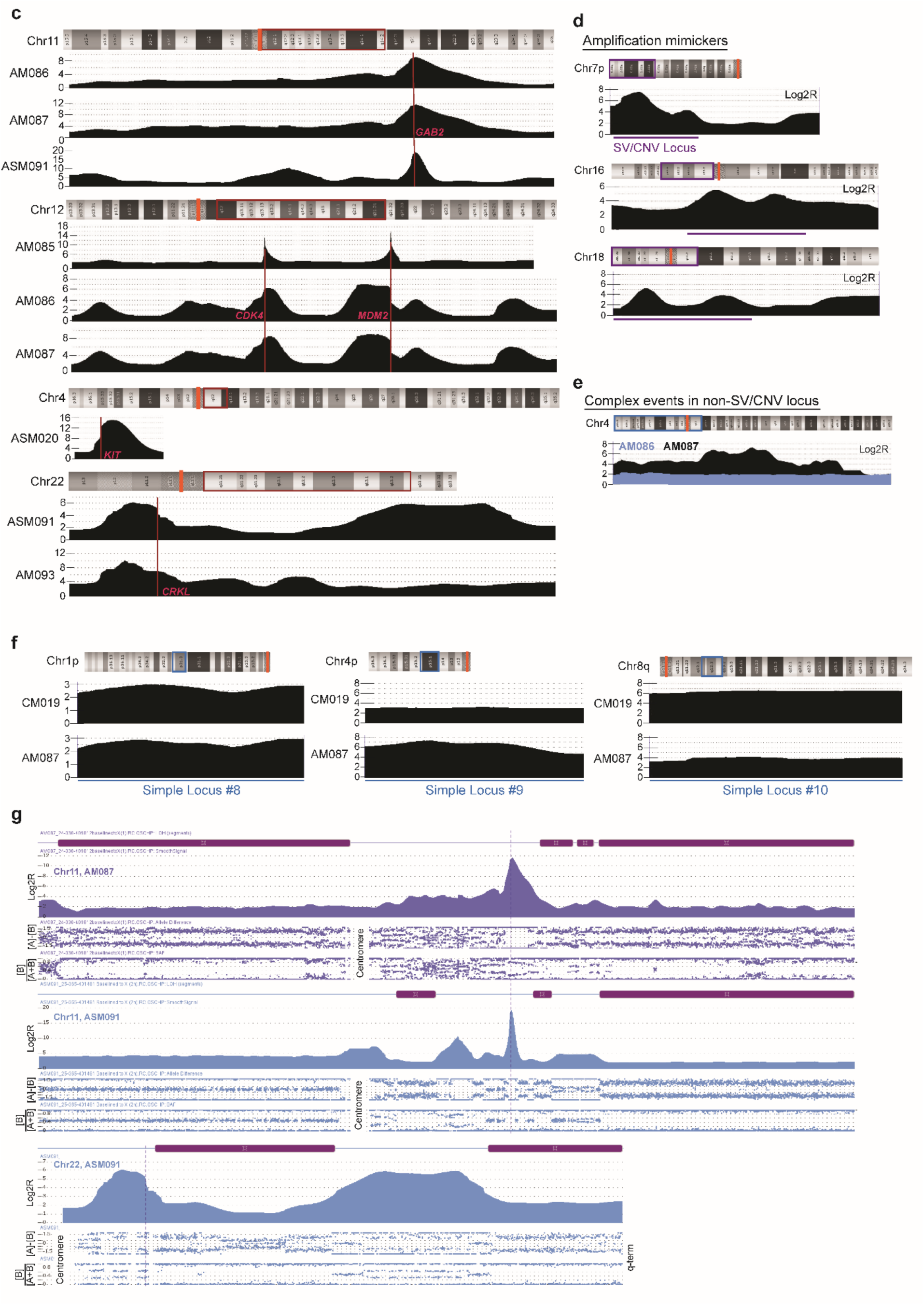
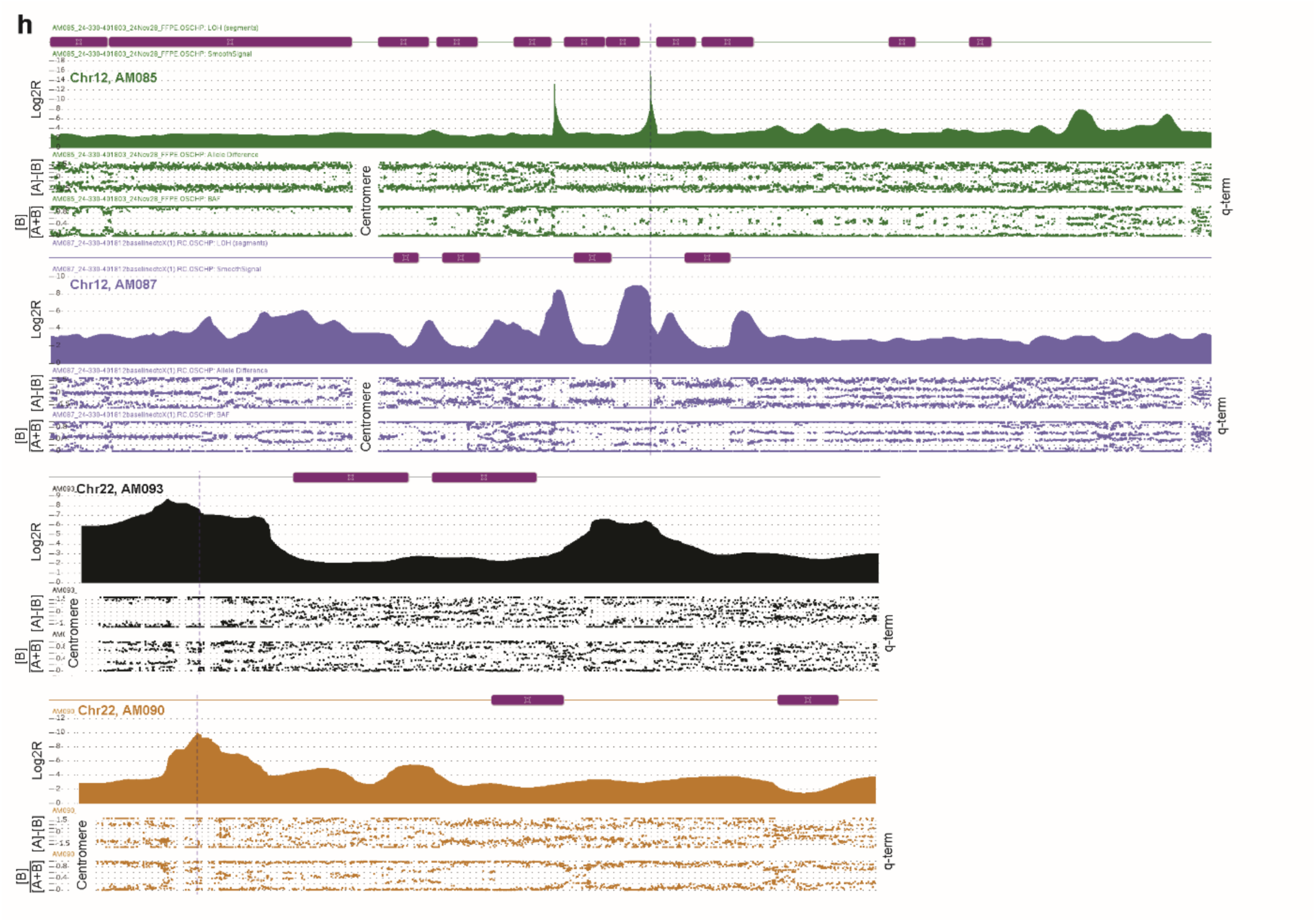
SV/CNV loci patterns by OncoScan CNV array. **a**, The top panel has a zoomed-in view of the CN change and SV connections between Chr11 and Chr12 by WGS as shown in **Extended Data Figure 6**. These regions are mapped to the OncoScan CNV array Log2R copy number tracing for the paired HCI-AM086 (blue) and HCI-AM087 (black) tumors from a male patient. Blue arrows point to where these peaks are represented on **b**, the CNV array whole genome view of HCI-AM087 from CNV array data (blue bars). The purple bar events are moderate copy number (CN) amplifications that required WGS support to be definitively called as SV/CNV loci. The top plot is the Log2R copy number tracing, middle plot is allele difference ratio ([A]-[B]), and bottom plot is B-allele frequency ([B]/([A]+[B])). Chromosomes are color-coded and labelled on the x-axis. **c**, All shown Log2R SV/CNV plots were initially identified by a high amplification spike on whole genome view as in **b**. The full chromosome is shown above the plot with centromere indicated by orange bar and viewed region by the colored box. The location of frequently amplified AM genes are shown by red vertical bars on Log2R traces. Note that these High-CN SV/CNV loci do not follow the gaussian patterns of double minute chromosomes or simple amplifications, and the SV/CNV loci comprise a small fraction of the chromosome arm. **d**, Moderate-CN SV/CNV loci in HCI-AM087 cannot be reliably distinguished from amplification events by CNV Array alone. **e**, One non-SV/CNV Simple Locus region that was not involved in the SV/CNV event by WGS underwent a distinct type of complex CNV change in HCI-AM087 (black profile), which was absent in the precursor HCI-AM086 tumor (blue profile). **f**, The non-SV/CNV Simple Loci used throughout the remainder of the report are shown here. **g-h**, Shown are full chromosome views of Log2R, allele difference and B-allele frequency for PDX models with High-CN SV/CNV loci. The purple rectangles represent LOH calls based on the allele ratio and B-allele frequencies. The centromere and q-terminus regions are labelled. Vertical dashed lines represent AM genes of interest: *GAB2* for Chr11, *MDM2* for Chr12, and *CRKL* for Chr22. LOH frequently marks the beginning and/or end of a SV/CNV locus. **g**, Several SV/CNV loci end in a terminal LOH that extends to the telomere. For Chr11 in HCI-AM087 the SV/CNV locus is highly amplified above the baseline whole chromosome LOH. **h**, HCI-AM085, HCI-AM090, and HCI-AM093 and HCI-AM087 Chr12 lack the terminal LOH seen in **g** but retain the LOH adjacent to SV/CNV loci.

**Extended Data Figure 9:**
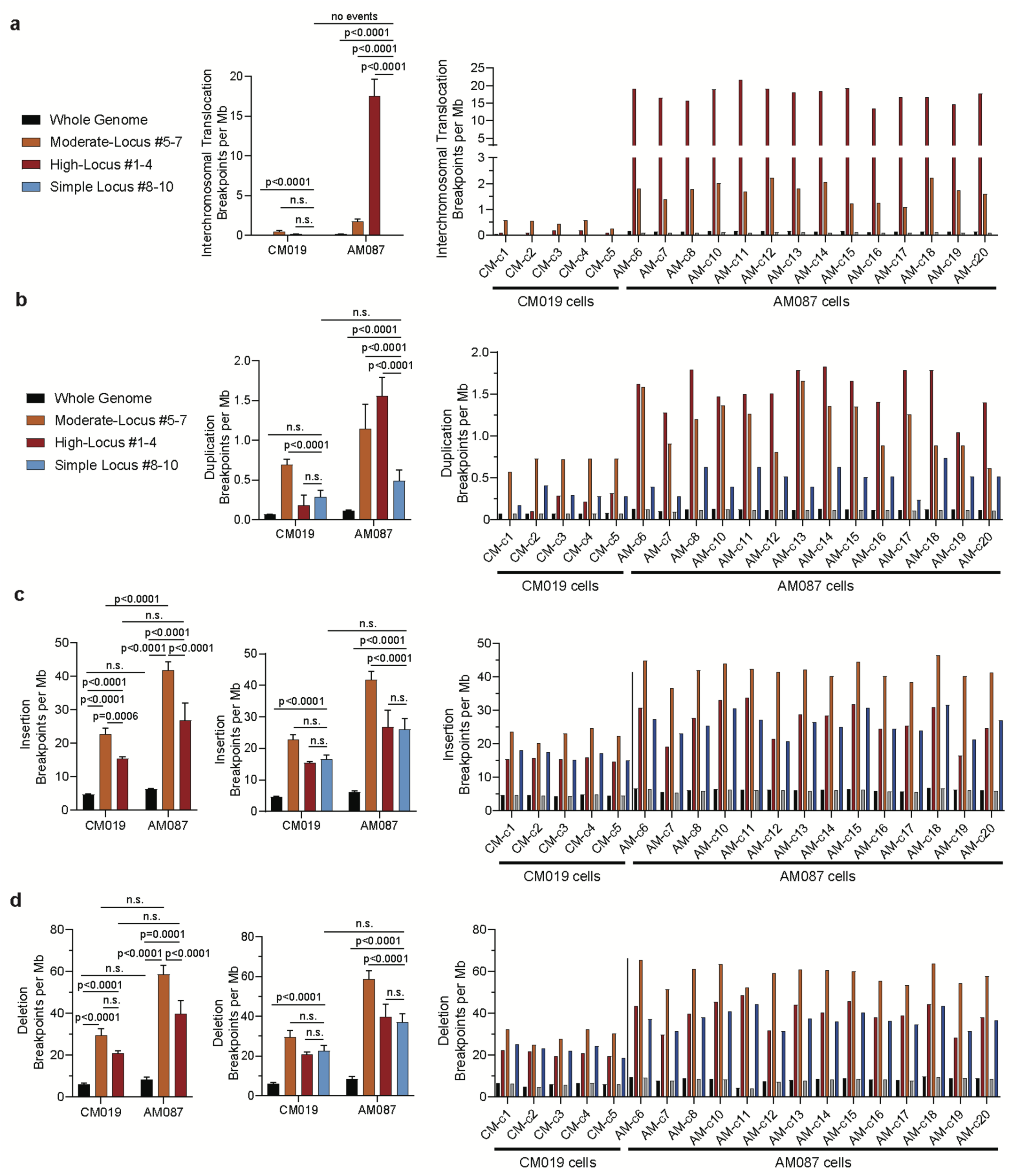
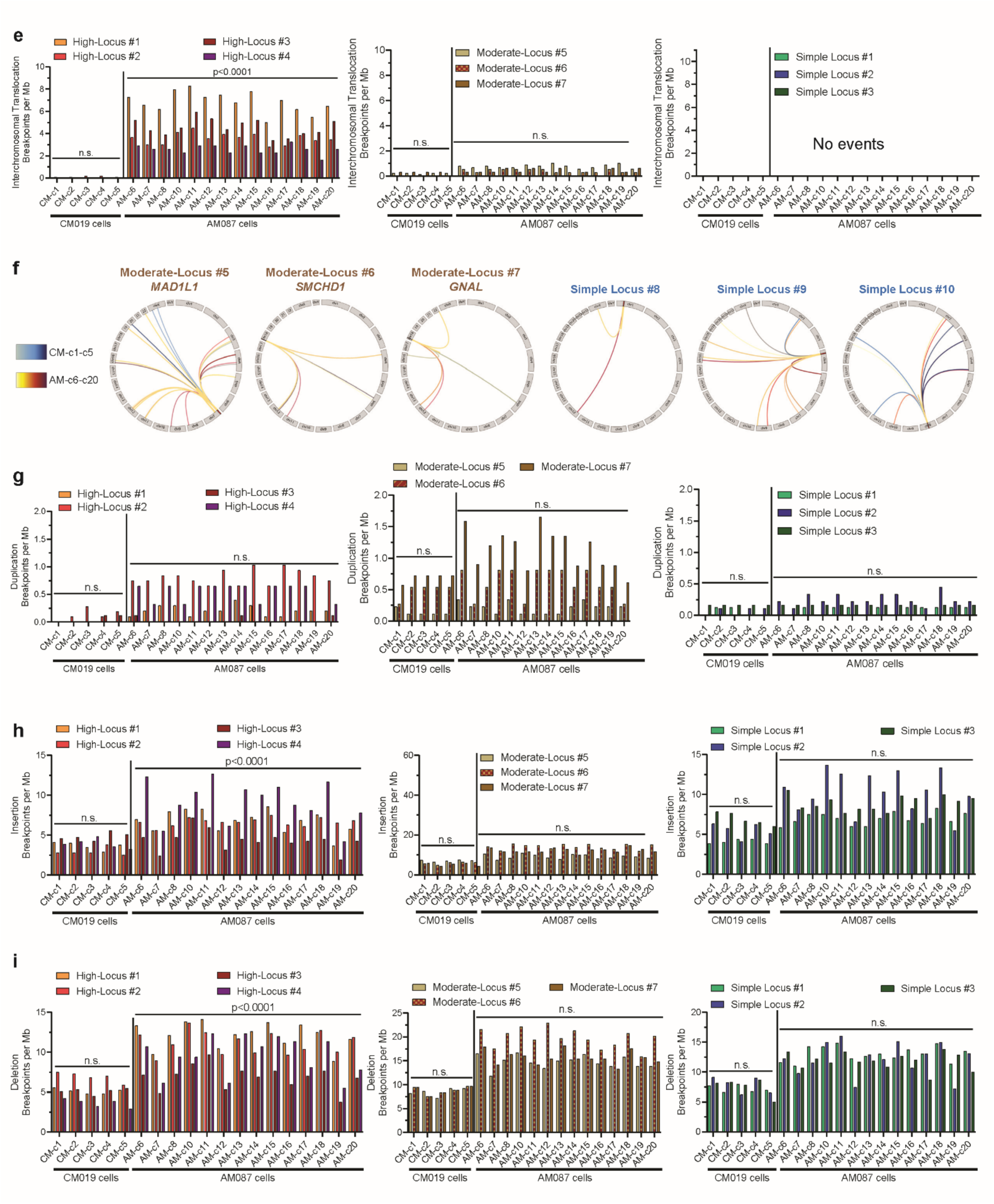
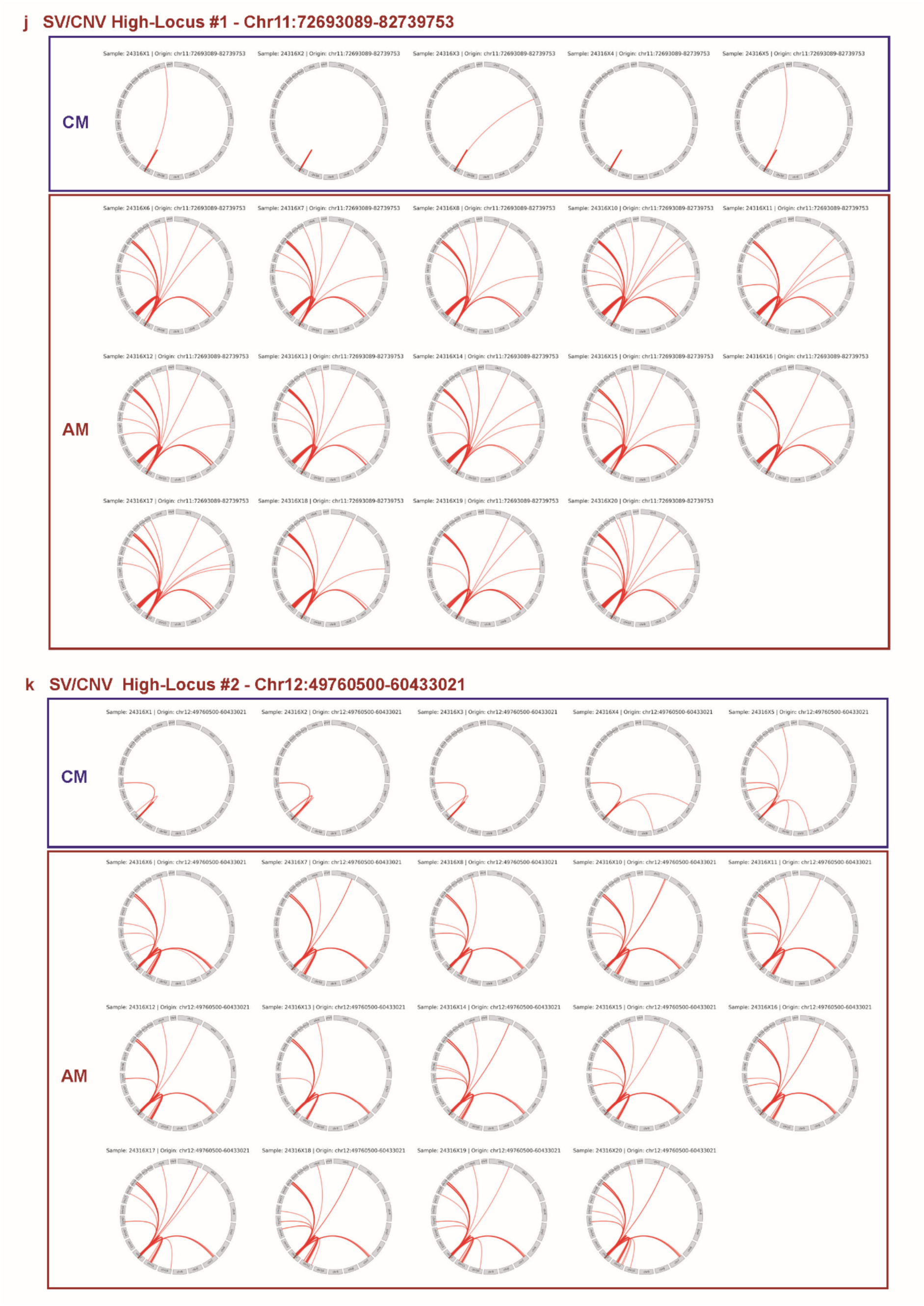
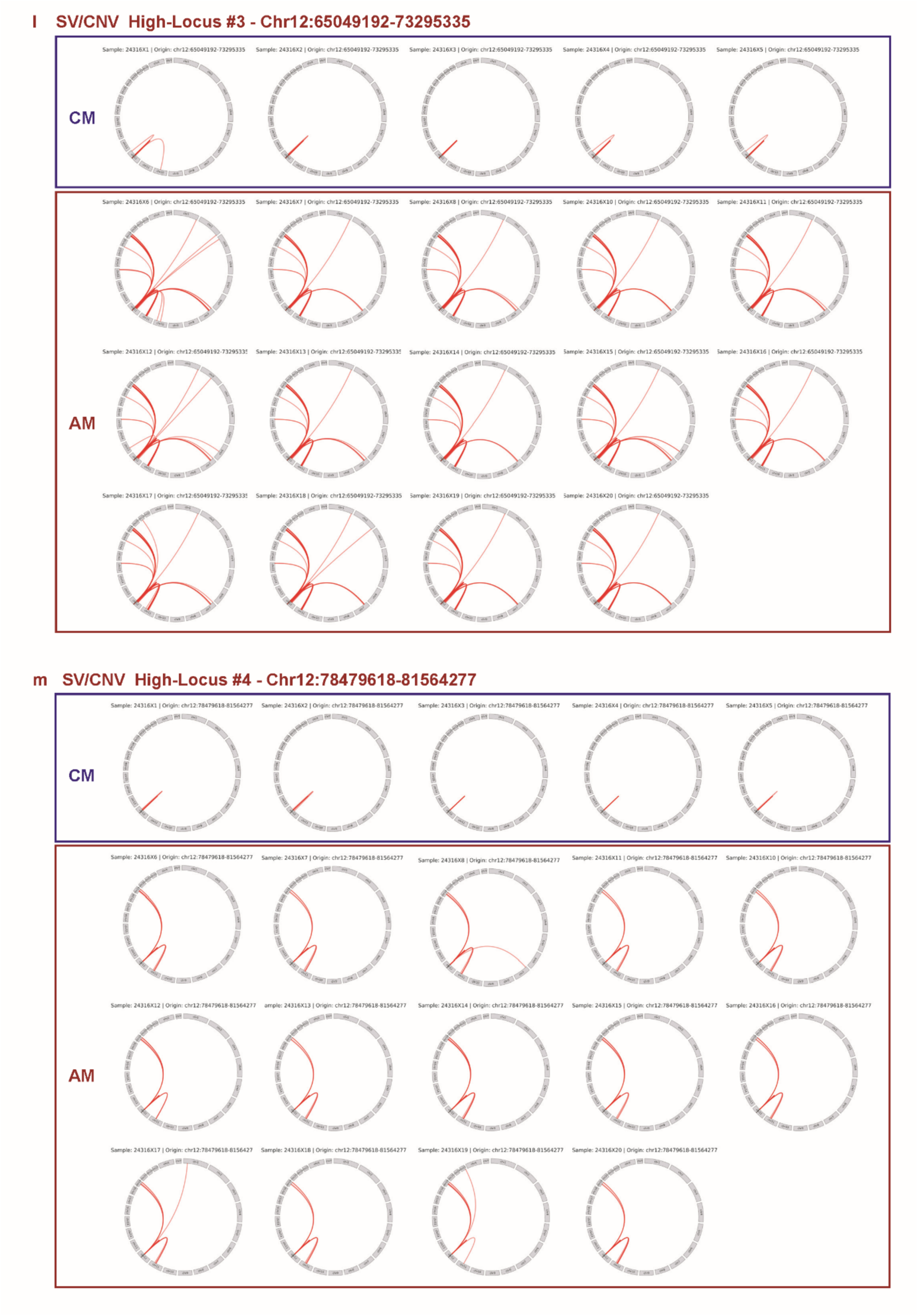
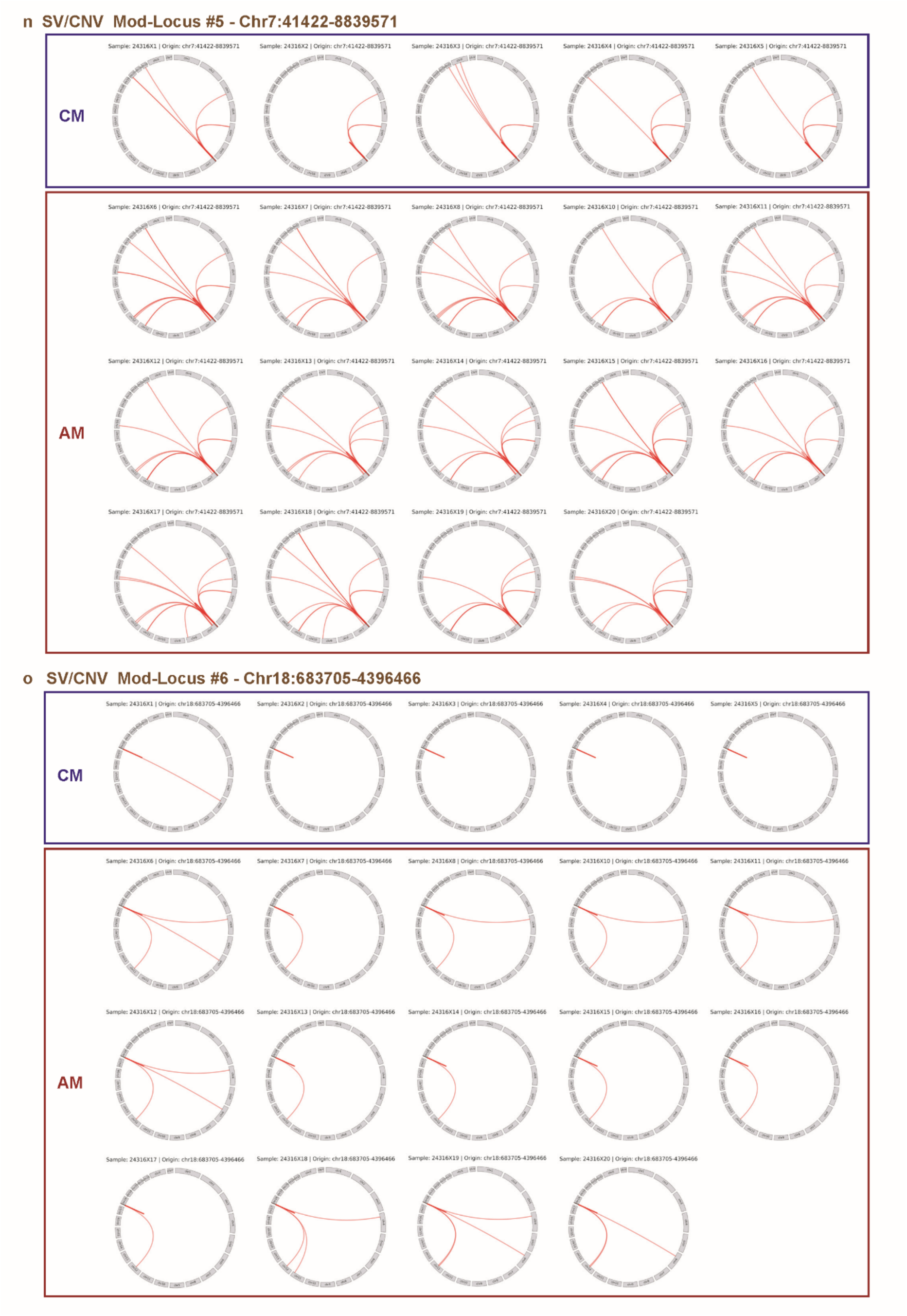
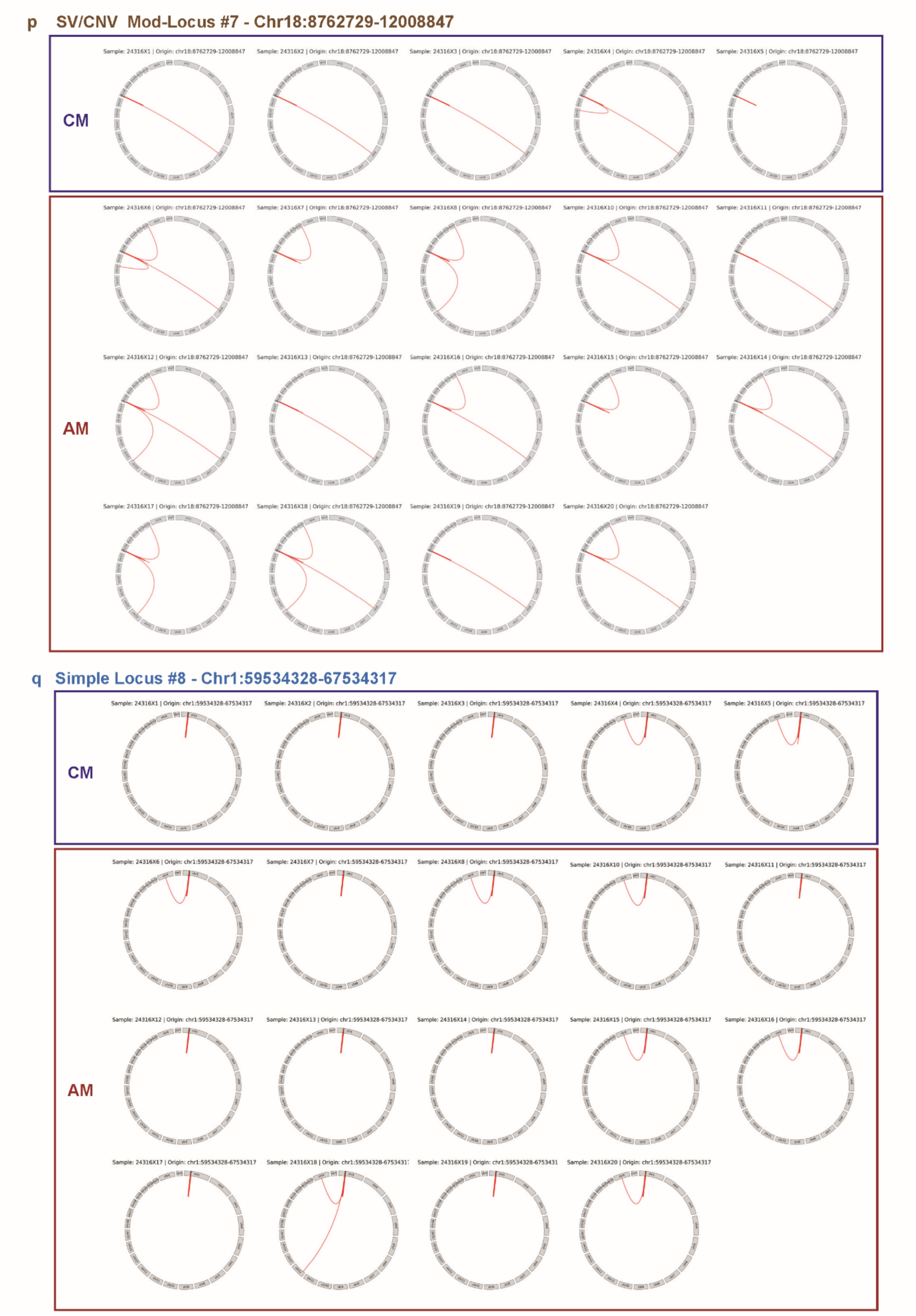
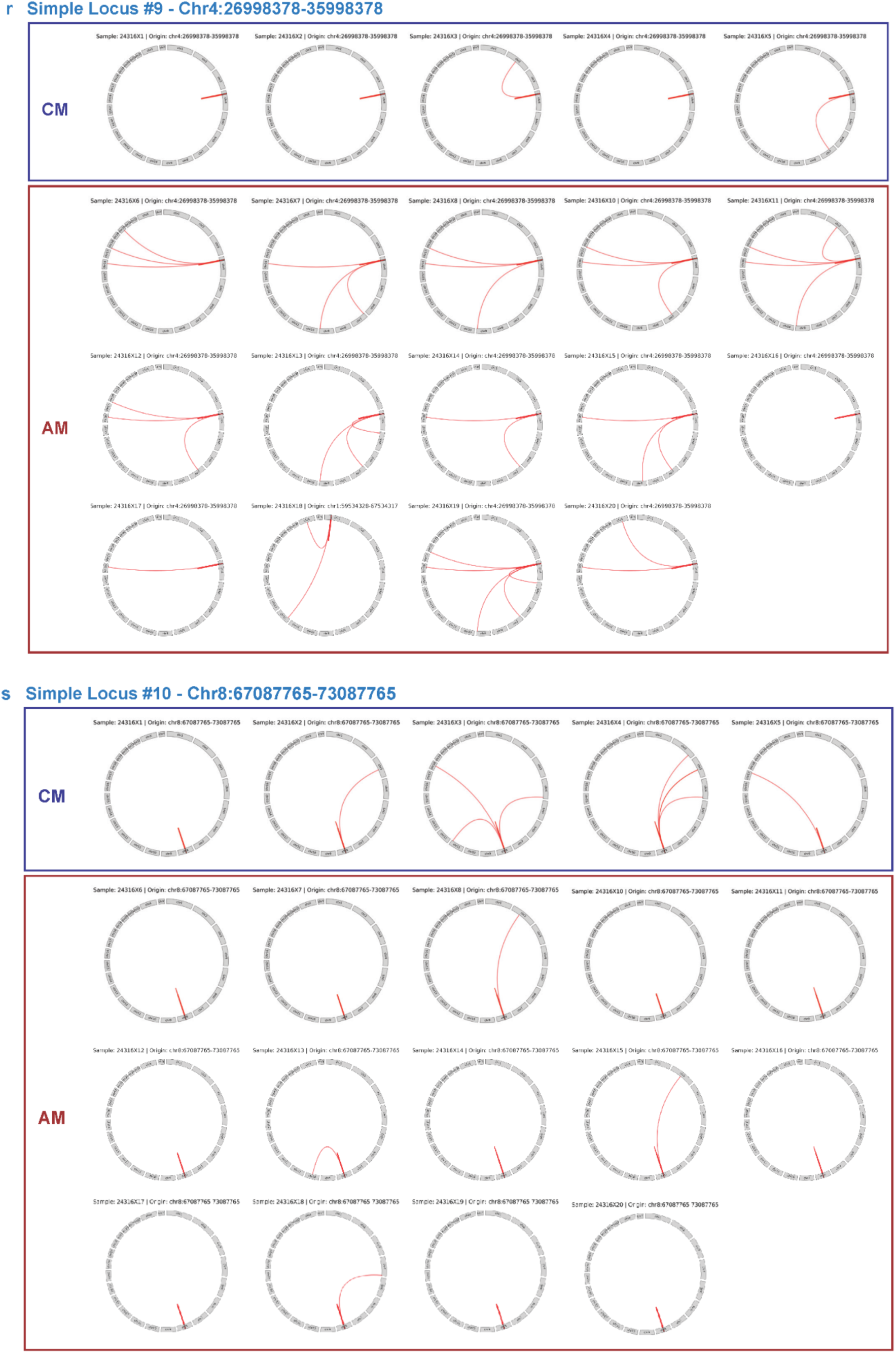
Translocations are enriched and highly variable in AM High-CN SV/CNV loci. **a-b**, The data from **Figure 3f** are replotted with the additional comparison to Simple Loci data. The right plots show breakpoint events in individual cells. **a**, Translocation events are greatly increased in HCI-AM087 SV/CNV loci compared to an absence at simple loci. **b**, There are double to triple the number of tandem duplication breakpoints in AM SV/CNV loci compared to background Simple Loci. For **c-d**, the whole genome, High-Loci #1-4, and Moderate-Loci #5-7 data are replotted in the left and middle graphs, with and without comparison to Simple Loci. The right graph shows breakpoint events in individual cells. **c**, Both CM and AM cells have increased insertion events at the Moderate-Loci compared to whole genome and Simple Loci, but High-Loci have equivocal insertion events to Simple Loci. Moderate-Loci have between 1.5x and 2x the number of insertions in AM compared to CM. **d**, Deletions are only moderately increased in Moderate-Loci compared to background Simple Loci, but are present at 1.5x to 2x the number in AM compared to CM. All datasets in **a-d** are demonstrated to have normal distribution by Shapiro-Wilk tests. Statistics are from paired t tests with a significance cutoff of p<0.001. **e**, Translocation events are significantly variable at High-Loci in AM only. **f**, Aggregate circos plots showing interchromosomal translocations in Moderate-Loci and Simple Loci. **g**, Tandem duplication events do not have significant variability between cells**. h-i**, Insertion and deletion breakpoints have significant variability only in High-Loci between AM cells. Significant variability in **e-i** was assessed by 2-way ANOVA. **j-s**, All interchromosomal translocations are shown for each cell at the seven SV/CNV loci of interest and the three Simple Loci. The circos plots are arranged in order of cell number order from left to right with locus name and hg38 coordinates.

**Extended Data Figure 10:**
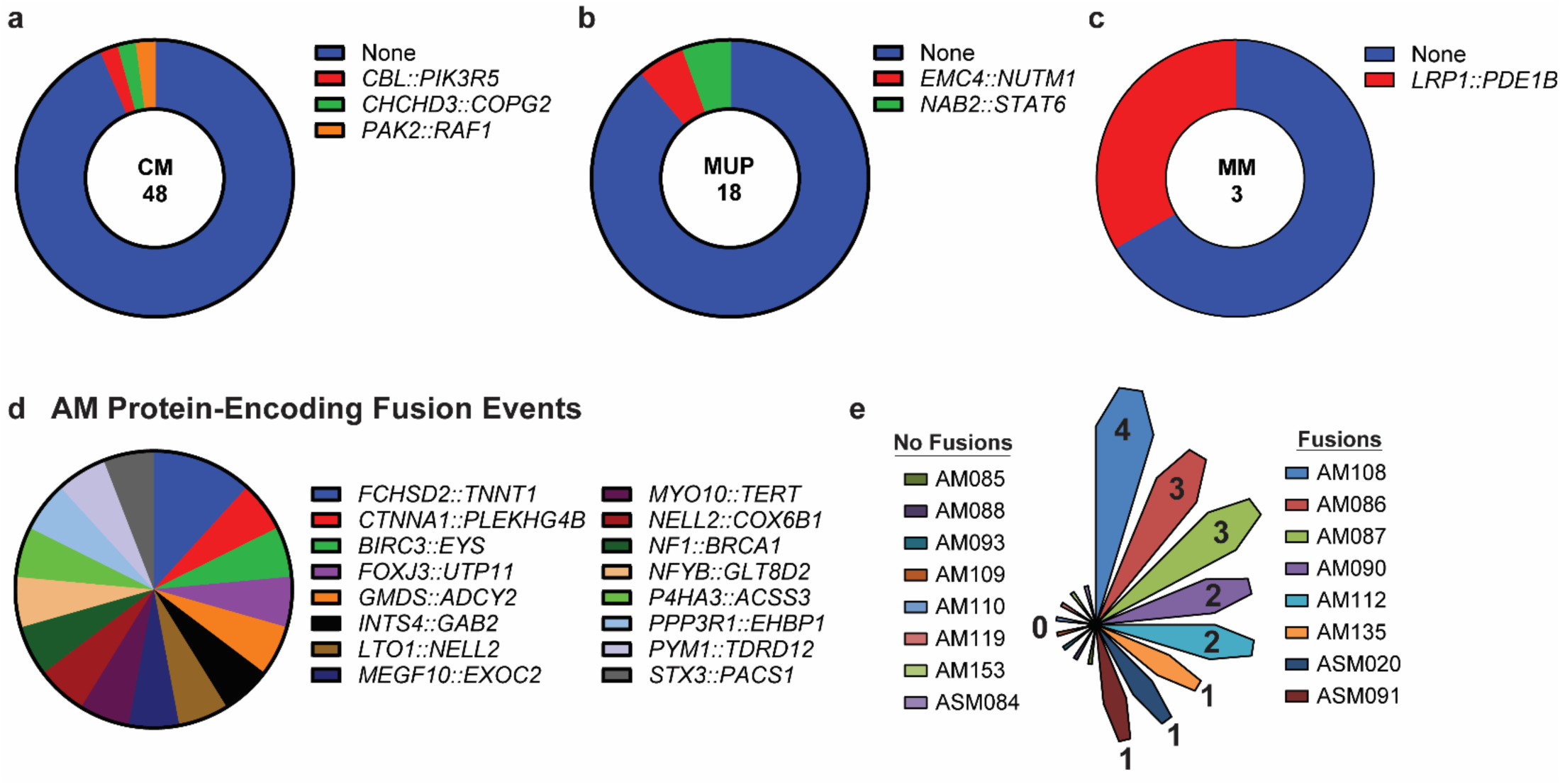
Protein-encoding fusions in AM. **a**, CM PDX had fusions in 8.3% of models (4/48) with two of the fusion models coming from the same patient. **b**, Fusions occurred in 11% (2/11) of MUP PDX models and in **c**, one of three mucosal melanoma PDX models. **d**, A subset of fusions in **Figure 5a** are predicted to encode for in-frame protein fusions. **e**, The number of predicted in-frame protein fusion products in each AM PDX model. Half of evaluated models have at least one predicted in-frame protein fusion product.

